# High-throughput smFRET analysis of freely diffusing nucleic acid molecules and associated proteins

**DOI:** 10.1101/651869

**Authors:** Maya Segal, Antonino Ingargiola, Eitan Lerner, Sang Yoon Chung, Jonathan A. White, Aaron Streets, S. Weiss, X. Michalet

**Author notes:** Email address:* (X. Michalet).

## Abstract

Single-molecule Förster resonance energy transfer (smFRET) is a powerful technique for nanometer-scale studies of single molecules. Solution-based smFRET, in particular, can be used to study equilibrium intra- and intermolecular conformations, binding/unbinding events and conformational changes under biologically relevant conditions without ensemble averaging. However, single-spot smFRET measurements in solution are slow. Here, we detail a high-throughput smFRET approach that extends the traditional single-spot confocal geometry to a multispot one. The excitation spots are optically conjugated to two custom silicon single photon avalanche diode (SPAD) arrays. Two-color excitation is implemented using a periodic acceptor excitation (PAX), allowing distinguishing between singly- and doubly-labeled molecules. We demonstrate the ability of this setup to rapidly and accurately determine FRET efficiencies and population stoichiometries by pooling the data collected independently from the multiple spots. We also show how the high throughput of this approach can be used to increase the temporal resolution of single-molecule FRET population characterization from minutes to seconds. Combined with microfluidics, this high-throughput approach will enable simple real-time kinetic studies as well as powerful molecular screening applications.

## 1. Introduction

Examining the three-dimensional structure of biomolecules is vital for understanding important biological functions. Techniques such as X-ray crystallography, nuclear magnetic resonance (NMR) spectroscopy, and cryogenic electron microscopy (cryo-EM) have been used in the past to determine biomolecular structures with nanometer spatial resolution. However, biomolecules are dynamic and undergo fluctuations that may not be captured by methods that require static samples. The result of these classical structural determination techniques is a static ‘snapshot’ of a dynamic process. While these high resolution ‘snapshots’ are hugely informative, they do not provide dynamic, temporal information of freely diffusing molecules in solution. In contrast, single molecule studies eliminate ensemble averaging and allow the possibility of capturing rare and transient conformational changes.

### 1.1. Background

Single-molecule Förster Resonance Energy Transfer (smFRET) techniques rely on the nanometer distance-dependence of the FRET efficiency between two spectrally matched dyes (the donor and the acceptor). This characteristic makes FRET a sensitive fluorescence-based molecular ruler that enables accurate determination of distances on the order of 3 – 10 nm [1]. Extension of this approach to the single-molecule level [2] has led to an ever growing number of applications, ranging from accurate measurement of equilibrium intra- and intermolecular conformations and binding/unbinding equilibria [3]. Combination with microfluidics [4, 5], electrokinetic trapping [6] or single-molecule manipulation techniques [7], later enabled studying conformational dynamics in solution at the single-molecule level. Recent developments have mainly focused on improving the reliability and resolution of distance measurements by smFRET [8, 9, 10], making it a useful complementary technique to X-ray crystallography and single-particle cryo-EM for exploring biomolecular structures. In particular, the ability of solution-based measurements to access molecular dynamics lays the foundation for time-resolved structure determination at the nanometer scale [3, 10].

### 1.2. HT-smFRET

Compared to measurements on immobilized molecules, solution-based measurements have the advantage of minimal perturbation of the studied molecule [11, 12, 13]. However, in order to ensure that only one molecule at a time traverses the excitation-detection volume, such that each transit can be clearly identified as a separate burst of photons, studies of single-molecules diffusing in solution are limited to low concentrations (≈ 100 pM or less). On one hand, this low concentration sensitivity makes smFRET a good tool for diagnostic applications in which patient samples are precious and target molecules may exist in very low abundance. On the other hand, the low concentration requirement poses challenges for the collection of large numbers of bursts needed for robust statistical analysis. In practice, this means that single-molecule measurements can last minutes to hours, limiting the application of smFRET to equilibrium reactions, unless combined with other techniques such as microfluidic mixers or some kind of trapping approach. Even then, accumulation of statistically significant data requires long acquisition times, due to the need of sequentially recording single-molecule data at each time point, such as in a mixer, or to sample enough individual molecule time-trajectories, as in the case of trapping. Parallel or multiplexed acquisition could overcome these challenges, without the need of, and possible artifacts associated with, immobilization, and expand smFRET applications to include fast, ultrasensitive clinical diagnostics and non-equilibrium kinetic studies.

Building on the recent development of single-photon avalanche diode (SPAD) arrays, we have recently demonstrated parallel detection of single-molecules and high-throughput smFRET (HT-smFRET) in solution by designing setups in which multiple excitation spots in the sample match the detector array pattern. Here, we provide details on our implementation as well as examples of applications, after a brief introduction of the SPAD array technology.

### 1.3. Custom silicon SPAD arrays vs CMOS SPAD arrays

Custom epitaxial silicon SPAD arrays used in this work were designed and fabricated by the SPAD lab at Politecnico di Milano (POLIMI, Milan, Italy) [14, 15, 16]. The detector modules include integrated active quenching circuits (iAQCs) designed to rapidly reset the SPADs in which an avalanche has been created upon absorption of an incoming photon. The modules are also equipped with timing electronics enabling single-photon counting, and, in some cases, with time-correlated single-photon counting electronics, enabling single-photon timing with ≈ 50 ps timing resolution [17]. It is worth noting that alternative SPAD array designs using standard complementary metal oxide semiconductor (CMOS) fabrication technology have also been developed during the past two decades [18]. While CMOS detectors afford larger scales (> 10^5^ SPADs versus < 10^3^ SPADs for the custom technology) ideal for wide-field imaging techniques, such as fluorescence lifetime imaging [19, 20] or high-throughput fluorescence correlation spectroscopy HT-FCS [21], it is our experience [22] that CMOS SPAD arrays still have a lower photon detection efficiency (PDE) and generally higher dark count rates (DCRs) than custom silicon SPAD detectors [14, 23] making them poor detectors for freely-diffusing single-molecule detection applications. Due to the fast pace of technological innovation in this field, this statement may become rapidly outdated. In addition to these fundamental differences, another important characteristic of custom technology SPADs is the possibility to manufacture larger individual SPADs while keeping low DCRs. This in turns simplifies precise optical alignment of the setup, making custom SPAD detectors ideal for single-molecule fluorescence studies.

This article is organized as follows: section 2 briefly describes the different multispot setups we have developed to emphasize common features and specificities. A detailed description of the 48-spot setup is provided in Appendix B. A brief outline of the analysis workflow for HT-smFRET data is presented in section 3, details being provided in Appendix C. Applications of HT-smFRET are discussed in section 4. We conclude by a discussion of on-going developments and future prospects for this technology.

## 2. Setup description

The general idea of a multispot setup involves replicating the usual confocal arrangement
of excitation spot and detector, with the constraint that each spot in the sample matches one
SPAD (further referred to as a “pixel”) in the SPAD array. There are multiple ways of achieving this goal, including using physical lenslet arrays, as we initially tried [24], or diffractive optical elements [25, 26]. The drawback of these approaches is the fixed spot pattern and possible aberrations thus obtained, which must be exactly matched to the fixed SPAD pattern in the emission path. This requires careful magnification adjustments, and cumbersome alignment steps, including adjusting a rotational degree of freedom. For these reasons, we chose a more flexible (if more expensive) solution using programmable liquid crystal on silicon spatial light modulators(LCOS-SLMs) [27, 28, 29, 30]. These devices can be used in direct space [31] or reciprocal space [32], as used in holography. As detailed below, this direct approach allows straightforward and real-time modification of the pattern and is capable of generating fairly uniform spots over the typical field of view of a high numerical aperture (NA) objective lens [33]. Alternatively, it is possible to use a line or sheet illumination pattern (Fig. 1B) and rely on out-of-focus light rejection by the geometry of the detector array itself, as we demonstrated with a linear array [17] and others have demonstrated with a 2D array [34] (although the latter demonstration was not a single-molecule experiment, the concept is applicable to smFRET). The drawback of these approaches, beside the increased background signal and inefficient excitation power distribution due to the absence of excitation light focusing, is the increased photobleaching resulting from the larger volume of sample in which fluorophore excitation takes place. This concern is diminished when using flowing samples, where exposure to excitation light is reduced by fast transit through the excitation region, as is the case when combining a multispot setup with microfluidics, as discussed later (Section 4.3.4).

**Figure 1:**
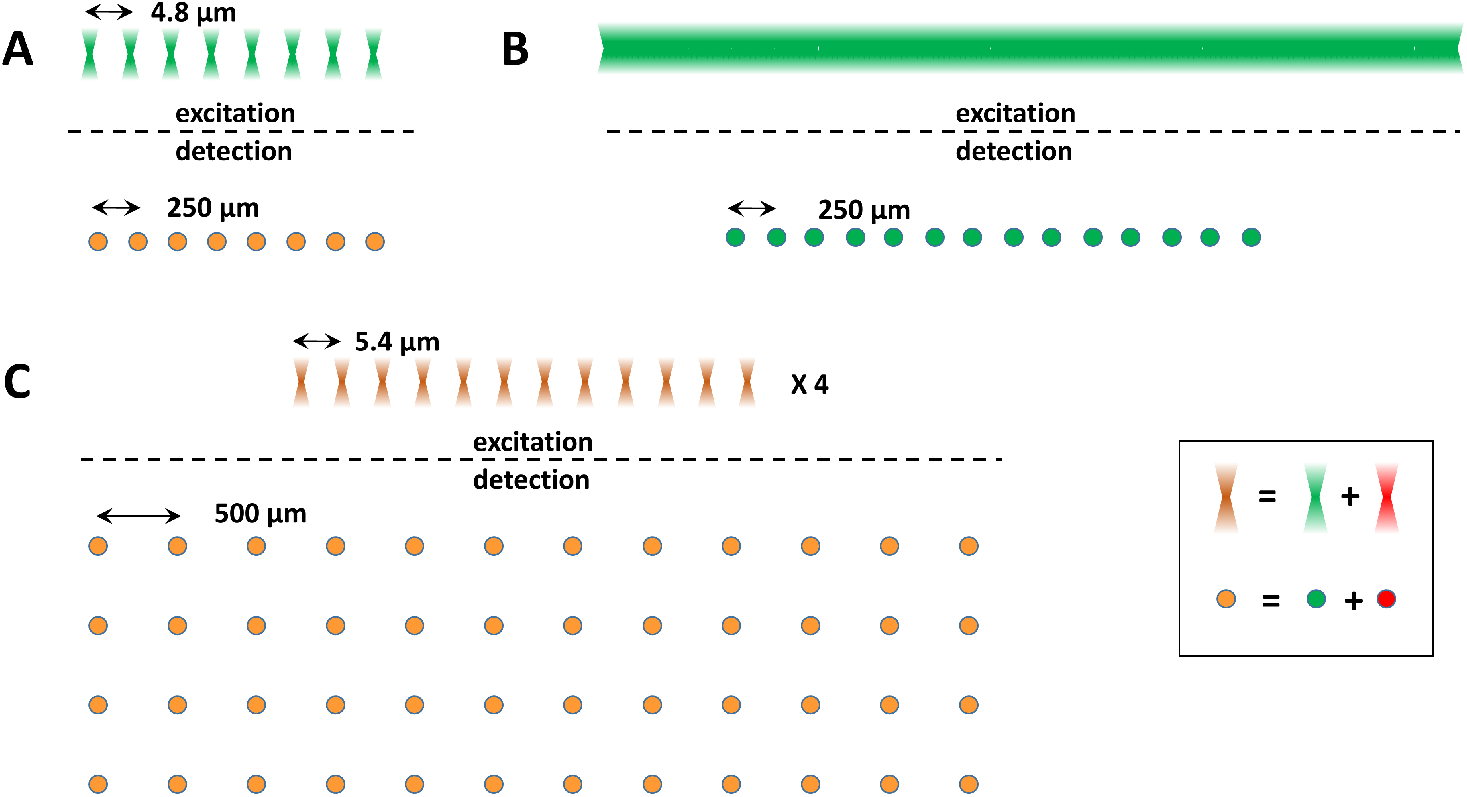
Different excitation and SPAD array geometries used in this work. A) Linear 8-spot and 8-SPAD array configuration. The 532 nm laser used in this setup was a high power (1 W) ps-pulsed laser (68 MHz). The setup was initially equipped with a single SPAD array (single color detection: green), and later upgraded with a second linear 8-SPAD array (red + green, represented in orange). The physical separation between excitation spots in the sample was 4.8 *μm* (top) [27], matching (after magnification) the 250 *μm* pitch of the SPADs in the array (SPAD diameter: 50 *μm*, bottom). B) A linear illumination pattern created with a cylindrical lens, using the same high power laser as in A was used to excite the fluorescence of samples. A linear 16-SPAD array (pitch: 250 *μm*, diameter: 50 *μm*) connected to a time-correlated (TCSPC) module was used to collect the emitted light from the conjugated spots in the sample [17]. C) Two patterns of 4×12 spots were generated in the sample by two high power (1 W) lasers (532 nm and 635 nm) and their associated LCOS-SLMs. The 5.4 *μm* distance between neighboring spots, matched, after magnification, the 500 *μm* distance between SPADs in the corresponding two 4×12 SPAD arrays (SPAD diameter: 50 *μm)* [30].

In our early efforts, we developed an 8-spot confocal microscope using a LCOS-SLM optically conjugated to a single linear 8-SPAD array. The 8-spot setup (Fig. 1A) employed a single continuous wave (CW) laser and was used to demonstrate HT-FCS and single-molecule detection [27]. We later added a second linear 8-SPAD array to the setup to enable two-color, 8-spot smFRET measurements [29]. Both setups used a 532 nm high-power 68 MHz pulsed laser for historical reasons, although we could not take advantage of the 8 ps pulsed laser excitation with these SPAD arrays. This configuration led to the development of a number of analysis tools allowing pooling of data acquired from separate spots for increased statistics (see section 3).

We next benchmarked a linear 32-SPAD array equipped with time-correlated single photon counting (TCSPC) readout electronics developed by POLIMI [35], using the same pulsed laser as before, but a simpler excitation optical train based on a cylindrical lens conjugated to the back focal plane of the microscope objective lens, in order to obtain a line illumination pattern, instead of an LCOS-SLM [17] (Fig. 1B). This test showed that time-resolved information (fluorescence lifetime decays) from multiple spots could be pooled together in order to speed up data acquisition, as already demonstrated for counting applications using CW excitation with the 8-spot setup.

After the development of larger SPAD arrays by our POLIMI collaborators, we upgraded our multispot smFRET setup with two 12×4 SPAD arrays [36, 30] (Fig. 1C). In addition to increasing the throughput, the 48-spot setup was designed with two CW lasers and two LCOS-SLMs for donor and acceptor dye excitation. Single-laser excitation is incapable of separating singly-labeled donor-only molecules (or doubly-labeled molecules with only one active acceptor dye) from molecules with low FRET efficiency (*E*, defined in section C), *i.e.* molecules in which the donor and acceptor inter-dye distance is greater than the Förster radius. Microsecond Alternated Laser EXcitation (μsALEX) using two laser excitations, was developed to overcome this challenge [37]. In μsALEX, two CW lasers are alternated on a time scale of a few tens of microseconds, shorter than the transit time of individual molecules through each excitation spot, allowing separation of doubly-labeled “FRET” species from singly-labeled donor-or acceptor-only molecules by calculating a simple uncorrected “stoichiometry” ratio (*S*, defined in section C). The combination of *E* and *S*, both calculated from single-burst intensities in each channel during each excitation period, enables “digital sorting” of different burst populations in the (*E, S*) plane, where all bursts detected during a measurement can be represented in a two-dimensional “ALEX histogram” and selected for further quantitative analyses [37, 38, 39, 40] (reviewed in [41]). Our setup uses a variant of this dual-excitation alternation scheme known as Periodic Acceptor EXcitation (PAX). PAX is a simplified implementation of ALEX where only the acceptor excitation is modulated and molecular sorting capabilities are preserved [42]. Comparing the performance of the 48-spot smFRET-PAX microscope to a standard single-spot μsALEX microscope, we found no difference in the quality of the data but a throughput increase approximately proportional to the number of SPADs, as expected. A schematic of the 48-spot smFRET-PAX setup is presented in Figure 12.

A setup incorporating two linear 32-SPAD arrays fabricated with a red-enhanced technology with better sensitivity [43], is currently under development for applications involving microfluidic mixers and will be described in a future publication.

The 48-spot setup is built with two 12×4 SPAD arrays and is equipped with two CW lasers with excitation wavelengths 532 nm (green) and 628 nm (red). A 12×4 lenslet array is generated using two LCOS-SLMs. In the 48-spot setup, only the acceptor laser (628 nm) is alternated. Setup details including the make and model of instrument parts for the 48-spot setup are included in Appendix B.

### 2.1. Excitation path optics

In PAX, the red laser is alternated by an acousto-optic modulator (AOM) and the green laser excitation is on continuously. Both laser beams are first expanded using a set of Keplerian telescopes, as shown in Figure 12A.3. Two periscopes raise the laser beams to a breadboard where the microscope body is placed. The laser beams are both expanded a second time (Figure 12A.4) in order to illuminate the LCOS-SLMs as uniformly as possible, as only phase modulation, not intensity modulation, is achievable with these devices.

Historically, PAX was introduced in our lab due to the availability of only one EOM (electro-optical modulator) to modulate both donor and acceptor lasers (unpublished). Later on, PAX was demonstrated using a modulatable red laser (acceptor excitation), the donor laser being non-modulatable [42]. Not using any AOM made this implementation of PAX simpler (and cheaper) than μsALEX, with no effect on data quality. In the smFRET-PAX setup described here, an AOM is used to alternate the red laser only. Thus, the main advantage of PAX over μsALEXis the simplified alignment, as only the red laser is diverted into the AOM. One disadvantage of PAX is additional photobleaching of fluorescent dyes due to the higher acceptor excitation power used to compensate for the lower detection efficiency in the red part of the spectrum. Further studies are needed to fully quantify this effect.

**Figure 2:**
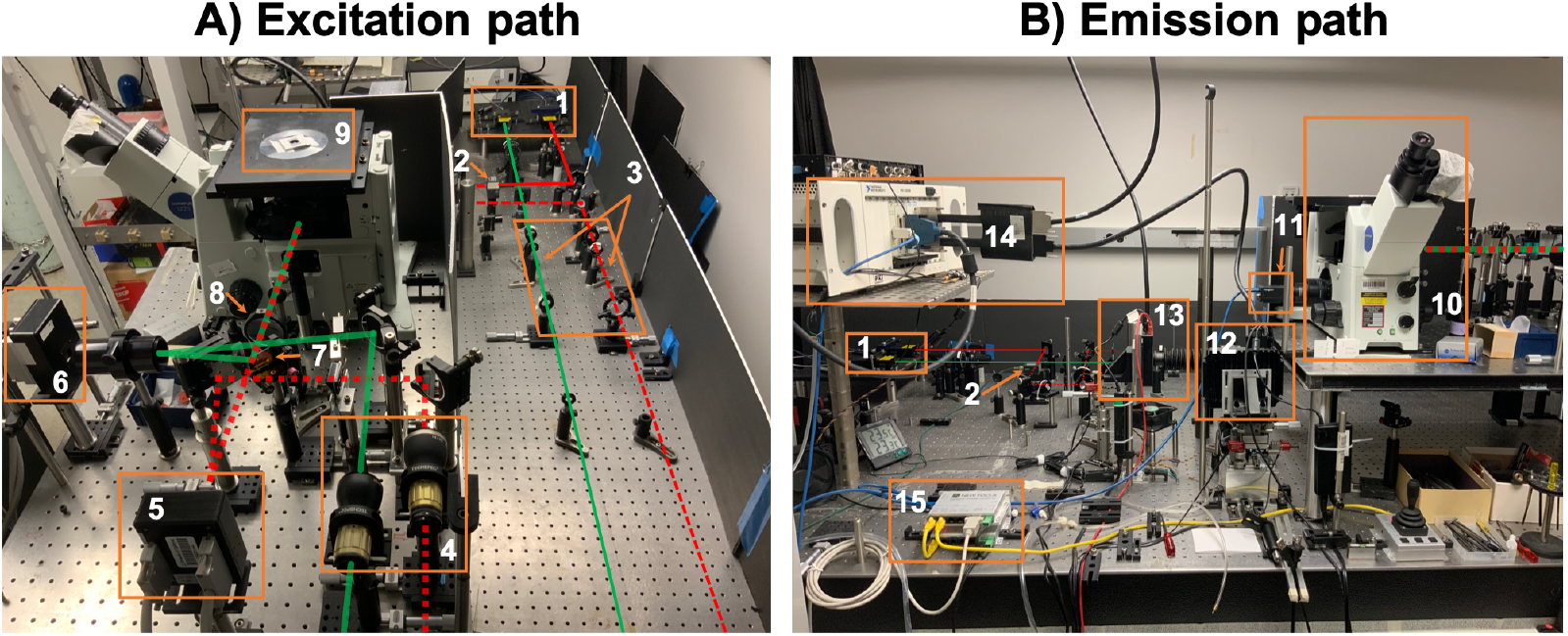
Photographs of the 48-spot setup. A) The excitation path consists of two 1 W CW lasers (1). Alternation of the red laser by the AOM (2) is indicated by red dashes. The lasers pass through a set of beam expanding lenses (3) followed by a second beam expansion (4) once on the upper breadboard. Both lasers are phase modulated by separate green (5) and red (6) LCOS-SLMs and the resulting beamlets are combined with a mixing dichroic (7) and recollimated by a recollimating lens (8) before entering the microscope body (10). B) Emission path optics showing the sCMOS camera (11) attached to the top side-port for alignment and the bottom path relaying the emitted fluorescence to the green (12) and red (13) SPAD arrays. Fluorescence emission is spectrally separated by a dichroic mirror and further filtered with emission filters (not visible), before being imaged onto two 12×4 SPAD arrays (12 and 13) mounted on micro-positioning stages powered by two micro-positioning drivers (15). The single-photon pulses from the SPAD arrays are sent to a programmable counting board (14) connected to the acquisition computer (not shown).

### 2.2. Phase modulation by LCOS-SLMs

In both 8-spot and 48-spot setups, the lenslet pattern is generated using LCOS-SLMs, where each spot is optically conjugated to a corresponding SPAD pixel. Patterns can be easily controlled using the relationship between the displacement and the phase delay of an incoming spherical wave, as detailed in the *LCOS Pattern Formation* section in the Supporting Information of ref. [29] and Appendix C in ref. [30].

Briefly, the LCOS-SLMs are programed to modulate the phase of the incident beams, effectively creating a 12×4 array of Fresnel microlenses matching the geometry of the two SPAD arrays. Spot generation by the LCOS-SLMs is obtained by sending two 8-bit encoded phase images (800×600 pixel), using the two LCOS-SLMs as “displays” attached to the host computer with a video card capable of supporting at least 3 displays. The phase images are generated with a custom LabVIEW program that computes the phase pattern using user inputs and supports automated pattern scanning as described in Ref. [30]. The `LCOS_LabVIEW` repository is available on GitHub (link).

The LCOS-SLMs each generate a 12×4 lenslet array, creating 48 separate excitation spots for each excitation wavelength. When properly aligned, the excitation spots overlap at the sample plane creating 48 dual-colored excitation spots. The lenslet array is focused at a user-specified focal length (see below for details) in front of the LCOS-SLM surface, as shown in Figure 13. The center and pitch of each pattern can be adjusted in the X and Y directions, and the pattern’s rotation can be changed using the `LCOS_LabVIEW` software (Figure 13, panels B and C). Demagnification in the excitation (83×), magnification in the emission (~ 60 × 1.5 = 90×) paths, and the geometry of the SPAD arrays dictate the pitch and resulting spot diameter:

- The spot pitch in the sample plane (5.4 μm) is defined by the pitch between adjacent SPADs (500 μm) divided by the emission path magnification (~ 90×).
- After demagnification by the excitation path, this translates into a 463 μm lenslet pitch on the LCOS-SLM, or 23.1 LCOS-SLM pixels. During alignment the pitch is optimized to account for the actual excitation path demagnification, by adjusting the pattern by fractions of LCOS-SLM pixels.

The focal lengths of the lenslet arrays are set to 36 mm and 30 mm for the green and red pattern respectively. The difference in the focal lengths accounts for the difference in PSF size for the 532 nm and 628 m wavelengths. Figure 3 shows the excitation pattern for the green and red lasers as visualized by a camera using a sample of high concentration ATTO 550 (panel A) and ATTO 647N (panel B) dyes. During alignment, the patterns are centered on the optical axis and their overlap is maximized. The overlap of the two excitations with respect to the optical axis is quantified by fitting the peak position and the Gaussian waist of each spot (Figure 3C). Details of the analysis are provided in the `pattern_profiling` alignment notebook (link) [30].

**Figure 3:**
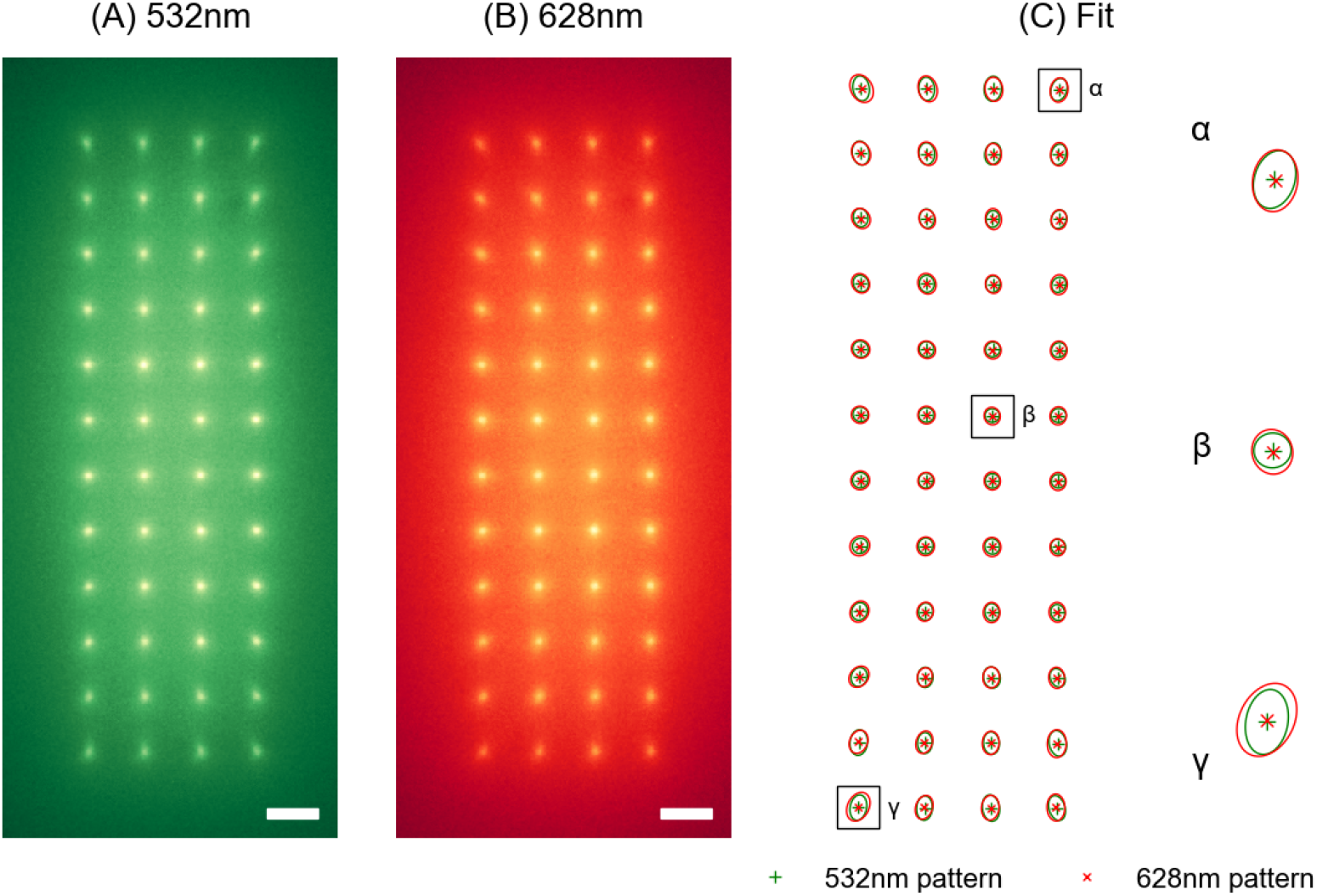
12×4 pattern generated by the two independent LCOS-SLMs. The elliptical shape and tilt of the Gaussian fits are due to residual optical aberrations. (A) Green (532 nm) 12×4 spot pattern. (B) Red (628 nm) 12×4 spot pattern. (C) To assess the alignment of the 12×4 patterns, each spot in the two images is fitted with a tilted 2D Gaussian function. The degree of overlap of green and red spots is determined by comparing the peak positions (cross and star) and the outline of the Gaussian waist (green and red ovals) of each green and red spot. Panels (A) and (B): images of a 100 nM mixture of ATTO 550 (green) and ATTO 647N (red) dyes, acquired separately with a CMOS camera installed on the microscope’s side port. Rightmost panel: *α,β*, and *γ* are close-ups of 3 representative spots in the 12×4 array. Scale bars = 5 μm. Reproduced from ref. [30].

#### 2.2.1. Background excitation reduction

To minimize background excitation, rectangular spatial filters (approximately 1 mm larger than the 12×4 pattern in both dimensions) are placed in front of the LCOS-SLMs, in order to block reflection by unused LCOS-SLM pixels. The phase modulated plane wave is reflected from the LCOS-SLMs creating the 12×4 excitation spot pattern. Unmodulated light from the pixels surrounding the pattern that is not blocked by the rectangular spatial filter is also reflected, creating specular reflections that contribute to the background signal. In order to suppress this residual specular reflection, a Bragg diffraction based beam-steering pattern is implemented around the lenslet array. The beam-steering pattern fills the region surrounding the 12×4 LCOS-SLM pattern with a periodic pattern that diffracts incoming stray light away from the back aperture of the objective (Figure 13A).

An example of the LabVIEW parameters for a 255 bit LCOS-SLM generating a 12×4 pattern for green and red excitations is represented in Figure 13B,C. The corresponding image of the 12×4 spot pattern formed at the sample plane is presented in Figure 3.

The software (LabVIEW & python) for generating the multispot LCOS-SLM pattern is freely available online (link), as part of the `multispot-software` repository used to align the LCOS-SLM pattern and SPAD arrays (link). During alignment, the acquisition software connects to the LCOS-SLM spot generation software. The positions of the LCOS-SLM patterns are scanned in two dimensions and the signal intensity from the center of the SPAD array is monitored. A detailed description of the procedure for aligning the 48-spot setup can be found in ref. [30].

### 2.3. Detection path

Fluorescence emission from the sample is collected by the same objective lens and passes through a dichroic mirror. The fluorescence emission is recollimated and sent through an emission dichroic mirror/filter cube where donor and acceptor emission wavelengths are separated before refocusing on their respective detectors. Each SPAD array is mounted on micro-positioners allowing adjustments of the detectors in all three directions. Adjustments in the transversal directions are performed with open loop piezo-motors controlled by software. The `picomoter` software used to control the micro-positioners is available as a GitHub repository (link). Alignment in the axial direction, being less critical, is done manually. The donor SPAD array is mounted on a rotation stage to fine-tune its orientation with respect to the acceptor SPAD array, allowing satisfactory overlap of the two 48-SPAD detectors.

### 2.4. SPAD arrays

The design and performance of the SPAD arrays have been described previously [16, 30]. Here, we briefly summarize this information.

#### 2.4.1. Dimensions and connectivity

The geometry of the two SPAD arrays consists of 12 rows of 4 pixels. Each SPAD pixel has and active area 50 *μ*m and is separated from its nearest neighbors by 500 *μ*m. The custom SPAD arrays fabricated by POLIMI are equipped with an internal field-programable gate array (FPGA) which can communicate with the acquisition PC via a USB 2 connection. Depending on the application, the FPGA firmware is used to merely report average counts per SPAD, or can send streams of individual photon timestamps to the host PC.

#### 2.4.2. Dark count rate and detection efficiency

The SPAD arrays are cooled to approximately −15°C in order to achieve the lowest possible DCR. The cooled SPADs have DCRs as low as 30 Hz, with an average of a few 100 Hz (donor channel: 531 ± 918H*z*, acceptor channel: 577 ± 1,261H*z*) [30]. A handful of SPADs have DCRs of a few kHz due to the difficulty of manufacturing large arrays with homogeneous performance. However, this noise level is adequate for smFRET studies where sample background is often comparable.

The detection efficiency of the standard technology SPAD arrays peaks at 550 nm, reaching a PDE of 45%. This makes it optimal for the detection of the donor dye (ATTO 550, emission peak: 576 nm), but less so for the acceptor dye (ATTO 647N, emission peak: 664 nm), for which the PDE decreases to 35% [36, 16, 23]. In particular, these values are 20 – 50% smaller than the most common SPAD detector used in single-spot smFRET measurements (SPCM-AQR, Excelitas Technology Corp., Waltham, MA) [33]. SPAD arrays fabricated with a red-enhanced technology with better sensitivity in the red region of the spectrum [44, 43] are currently being evaluated in our laboratory, and will reduce the performance gap between donor and acceptor detection efficiency.

#### 2.4.3. Afterpulsing

Like single SPADs, SPAD arrays experience afterpulsing due to the non-zero trapping probability of carriers created during an avalanche and late release after the initial counting event, resulting in spurious counts. The typical time scale of these delayed signals depends on the device and can range from hundreds of nanoseconds to several microseconds, resulting in noticeable autocorrelation function (ACF) amplitude when performing FCS analysis [27, 29]. While there are techniques to correct for this effect [45], they require good separation between the time scale of afterpulsing and that of the phenomenon of interest. Some of the SPAD arrays we have tested do not satisfy this condition, making it challenging to reliably extract short time scale (< 1 − 10 *μ*s) parameters by ACF analysis only, although the contribution of afterpulsing can otherwise be accounted fairly well using a power law fit [27, 29]. Instead, short time-scale correlation analysis can be accomplished via cross-correlation function (CCF) analysis if the signal is split equally between two different detectors [46], but this requires twice as many SPAD arrays.

Provided that detector deadtime and afterpulsing effects are independent [47], the afterpulsing probability, *P_a_*, can be estimated simply, by recording counts under constant illumination:

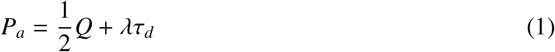

where Q is the Mandel parameter (Eq. 2) characterizing the recorded signal, *S. λ* is the incident count rate and *τ_d_* is the deadtime (120 ns in the SPAD arrays discussed here).

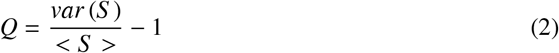

For a constant illumination where 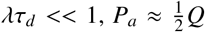, which is in general small (for a pure Poisson process, *Q* = 0, therefore *P_a_* is a measure of the departure from this ideal situation). The measured afterpulsing probability (several percents) is currently higher in SPAD arrays than in single SPADs where *P_a_* < 0.1% [29], but this will most certainly be improved in future generations of detectors.

#### 2.4.4. Crosstalk

Another important specificity of SPAD arrays is the potential occurrence of electrical and optical crosstalk effects. Electrical crosstalk is due to parasitic signals generated in the compact circuitry surrounding the detectors, and can in principle be eliminated with careful design. On the other hand, optical crosstalk is due to emission of secondary photons during the avalanche [48] and is independent of the type of setup the detector is used in [49, 50]. These secondary photons can propagate to neighboring or distant pixels and trigger avalanches in them [51]. The resulting spurious signals occur at very short time scales, set by the avalanche quenching time (< 20 ns for SPADs equipped with iAQCs [52]). Crosstalk percentage can be estimated by a simple dark count measurement, and analyzed by CCF or mere counting [53, 29, 54]. Defining *C_c_* as the number of coincident counts in two pixels A and B in a time window Δ*T* slightly larger than the crosstalk time scale, the crosstalk probability, *P_c_*, can be estimated from the number of counts in SPAD A and B, *N_A_* and *N_B_*, as:

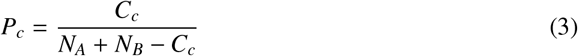

In a recent study, we thoroughly characterized the magnitude of optical crosstalk in our 48-SPAD arrays [54] and found it to be of the order of 1. 1 × 10^−3^ for nearest-neighbors and 1.5× 10^−4^ for nearest-diagonal pixels. The crosstalk probability for pixels further apart drops to even more negligible levels for these newer SPAD arrays, demonstrating a significant improvement over previous models [29]. The improved optical crosstalk probability is attributed to the high doping levels (> 2 × 10^−19^cm^−3^) used in the new fabrication process, which reduces propagation of photons through the silicon layer eliminating reflections from the bottom of the chip [48].

Yet another potential source of optical crosstalk can come from the physical proximity of the volumes sampled by nearby pixels: in diffraction-limited setups, molecules excited at and emitting from spot *n* must have their signal collected and imaged by pixel *n* only, in each channel. In an ideal setup, the image of each excitation/detection spot is a point-spread-function (PSF) whose extension should be limited to a single pixel, and in particular, should not overlap neighboring pixels. The SPAD arrays we use have a pitch-to-diameter ratio of 500 *μm*/50 *μm* = 10, and the detection path magnification (*M* = 90) is such that the full-width of the PSF’s image (≈ *Mλ*) is comparable to the SPAD diameter, ensuring no overlap between the PSF image of neighboring spots.

Note that at this time, there are no commercially available custom SPAD arrays, although we hope this technology will become available in the near future. CMOS SPAD arrays are available, however single-molecule detection is not practical with these devices.

### 2.5. Multispot data acquisition

A n-SPAD array output consists in *n* independent streams of “pulses”, each pulse corresponding to an avalanche due to one of several kinds of events: photon detection, afterpulse, crosstalk pulse, or dark count. These electric pulses are generally shaped by onboard electronics (transistor-transistor logic (TTL) or nuclear instrumentation module (NIM) pulses are standard) and readout by internal or external processing electronics. The POLIMI detectors we have used were characterized by a variety of output signal configurations:

- independent TTL signals with one Bayonet Neill–Concelman (BNC) cable per channel for the 8-SPAD arrays [27, 28, 29],
- Low-voltage differential signaling (LVDS) converted to TTL signals by an external board [30], and finally,
- independent fast-timing signals and counting signals [17].

The latter two detector modules incorporate an FPGA for signal conditioning (resulting in the TTL or LVDS pulses mentioned previously), and if needed, actual photon counting. Data processed by the FPGA has a 50 ns resolution time-stamp and pixel identification for each count and can be transferred asynchronously via USB connection to the host PC, which makes these devices particularly easy to use. In the case of TCSPC measurements [17], the fast timing signals were fed to a separate module incorporating time-to-amplitude converters (TACs) connected to the laser trigger. The TAC outputs, converted to nanotiming information, and combined with channel identification and macrotiming information provided by the clock of an embedded FPGA, were transferred asynchronously via USB connection to the host PC [35].

However, when two separate detectors are used simultaneously, as needed for FRET measurements, synchronization of the two series of photon streams originating from both detectors requires that all events be processed using a common clock. As this synchronization has not yet been implemented, we resorted to a different approach, feeding pulses from both detectors to a single, external counting board.

The counting board used in all works cited previously (except the TCSPC work) is programmable and allows buffered asynchronous transfer of data to the host computer (PXI-7813R, National Instruments, Austin, TX). Supporting up to 160 TTL inputs, it is in principle sufficient to handle up to three 48-SPAD arrays. Data consists of a 12.5 ns resolution, 24-bit time-stamp for each photon, as well as a 7-bit pixel number. The theoretical throughput of the PXI-7813R is 40 MHz, but sustained transfer rates are generally lower, which can result in lost counts at high count rates. Fortunately, this is not an issue in smFRET, where the average count rate per channel is rarely larger than 10 kHz, and while instantaneous peak count rates are on the order of a few MHz per pixel (see below), each pixel is uncorrelated to the others. The LabVIEW FPGA code for the counting board we used is available in the `Multichannel-Timestamper` repository (link).

For PAX measurements, an additional board (PXI-6602, National Instruments), whose base clock is fed to the programmable board, is used to generate the digital modulation signal sent to the AOM. This synchronization is critical to be able to assign each recorded photon to one of the two excitation periods of each alternation.

### 2.6. Multispot data saving

Data recorded during these experiments is processed in real time and displayed as binned time traces, or when dealing with large number of channels, as color-coded binned intensity charts, in order to monitor the experiment. Simultaneously, the data comprised of a timestamp and SPAD ID number for each photon is streamed to disk as a binary file. In order to facilitate handling of the different configurations of pixel number and data types (counting or TCSPC), this raw binary data is next converted with the addition of user-provided metadata stored in a YAML file into a general and open source photon-counting data file format, (Photon-HDF5) [55, 56], using the `phconvert` python library (link). This file format was designed for maximum flexibility and storage efficiency, and can be easily used with most programming languages. Because it is extensively documented, and compliant files contain all the information necessary to interpret and analyze single or multispot data, we hope it is a tool that the community of diffusing single-molecule spectroscopists will use. In particular, it is accompanied by `phconvert`, a tool that allows conversion of several commercial file formats into Photon-HDF5 files, which will facilitate data sharing and analysis cross-validation.

In our workflow, conversion from proprietary binary file to Photon-HDF5 is performed as soon as the binary file is saved, using a second computer that monitors the data folder. This conversion can be followed by scripted smFRET analysis as described below, freeing the data acquisition computer for further acquisition, as needed in high-throughput applications. With the advent of fast solid state drives (SSD) and increasing number of central processing unit (CPU) cores, it is likely that this division of tasks will not be needed in the future, allowing real-time data analysis and display on a single computer.

## 3. Data analysis

In this section, we present a brief overview of the typical workflow, with special emphasis on the multispot specific steps. Notations and definitions, as well as details on the analysis can be found in Appendix C and in ref. [29, 30].

### 3.1. smFRET burst analysis

smFRET analysis of freely diffusing molecules in solution involves many steps, the basis of which has been discussed in many publications (e.g. [57, 58, 59, 38, 60, 61, 62, 29]), however the very complexity of this type of analysis makes the results sensitive to many details (such as parameter values). For instance, in order to be able to compare methods when a new approach is introduced or when a result raises questions, it is important to have access to not only raw data sets but also analysis parameters, steps performed during the analysis, and implementation details.

While it is not the purpose of this article to discuss the implications of these requirements in depth, the best way to guarantee reproducibility and testability is to provide a detailed record of the analysis, including inputs and outputs, as well as the complete list of analysis steps. This is best achieved by providing the source code (e.g. [63, 64]), but also requires documentation of both code and workflow. In this work, we mostly use FRETBursts, an open-source and fully documented python package available at https://github.com/tritemio/FRETBursts, allowing reproducible single-molecule burst analysis [63]. Data analysis steps and results are recorded within Jupyter notebooks, linked to in the different figure captions or throughout the text of this article. In addition, ALiX, a free, standalone LabVIEW executable performing essentially the same functions [29], was used and is available at https://sites.google.com/a/g.ucla.edu/alix/. Logbooks generated during the analysis are provided as Supporting Information. While ALiX’s source code is not released yet, mostly because it is developed with the graphical language Lab-VIEW, for which no simple “reader” exists, it is available upon request from the authors, and is developed under version control for traceability. An extensive online manual is also available (link).

To our knowledge, both packages are the only ones to support multispot analysis.

### 3.2. smFRET multispot burst analysis

Multispot analysis can be divided into three different types:

1. Independent single-spot analysis
2. Pooled multispot data analysis
3. Spot correlation analysis

In independent single-spot analysis each spot is treated as a separate measurement. This type of analysis is appropriate for geometries in which each spot probes a different sample, such as parallel microfluidic channels probed by one spot each.

The second case involves data collection from each spot, independent burst analysis for the different data sets, and pooling of burst data from all spots to increase statistics.

Finally, in the third case, data from different spots can be correlated, for instance, using intensity CCF analysis, in order to obtain transport coefficients or any other type of information unobtainable from individual spot analysis. This type of analysis is used for crosstalk estimation (see section 2.4.4) and is illustrated in the microfluidic section of this article.

Many factors must be considered in order to implement robust pooled smFRET analysis. Indeed, a measurement performed with an N-spot setup is actually similar to N distinct measurements performed simultaneously on the same sample. Differences between these individual recordings are due to small differences in the characteristics of each excitation/detection volume (including peak intensity), as well as in the performance of each SPAD (in particular DCR and afterpulsing). Due to the independent alignment of each illumination pattern and each detector, these differences are amplified by the number of excitation lasers and the number of detection channels, underscoring the importance of a good alignment procedure and thermal and mechanical stability.

In order to pool burst data from each spot into a single global data set, it is necessary to quantify these differences and determine the relevant correction factors (these correction factors are discussed in Appendix C.8. Correction factors involved in FCS analysis were discussed in [27]). We have illustrated this procedure in ref. [30] and summarize the results in the first part of the next section, which describes examples of HT-smFRET analysis.

Details on smFRET-PAX analysis can be found in Appendix C and in the *48-spot-smFRET-PAX-analysis* repository (link).

## 4. Applications of HT-smFRET

### 4.1. Equilibrium smFRET measurements

#### 4.1.1. HT-smFRET-PAX of freely diffusing dsDNA

In order to demonstrate the increased throughput of the 48-spot smFRET-PAX setup, we first performed measurements of doubly-labeled 40-base pair (bp) double-stranded DNA (dsDNA) molecules. ATTO 550 (donor dye) and ATTO 647N (acceptor dye), were each attached to a single strand, different samples being characterized by different interdye distances, as detailed in ref. [29]. Here, we limit ourselves to interdye distances of 12 bp and 22 bp [30]. Measurements were performed on dilute samples (≈ 100 pM) in TE 50 buffer, a minimal DNA storage buffer containing 10 mM Tris pH = 8.0, 1 mM EDTA, and 50 mM NaCl. Each sample was measured on a standard single-spot μsALEX setup, followed by measurement on the 48-spot PAX setup. This ensured that sample conditions were identical for comparison of setup characteristics.

#### 4.1.2. Burst search and selection

As described in Appendix C, smFRET analysis is performed in three steps:

1. Background estimation
2. Burst search
3. Burst selection

To account for possible fluctuations in background levels, background rate estimation for each photon stream at each spot was preformed using a 10 s sliding time window. Burst search was then performed for each spot using a standard sliding window algorithm, defining the local total count rate using *m* = 10 consecutive photons, and using a constant threshold to define burst start and end (50 × 10^3^ cps or 50 kHz). After burst search, different selection criteria can be applied to further isolate burst subpopulations. Typically, a first burst selection based on a minimum background-corrected total count (e.g. ≥ 30 photons) is used to only keep bursts with sufficient signal-to-noise ratio. Further selections can be applied to separate FRET species from donor-only and acceptor-only molecules. This can be achieved by selecting bursts whose total acceptor signal during both donor excitation and acceptor excitation, 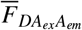, is larger than a minimum value, see Figure 4. However, it is generally simpler to use the 2-dimensional (2-D) *E* – *S* histogram discussed next to identify and graphically select sub-populations.

**Figure 4:**
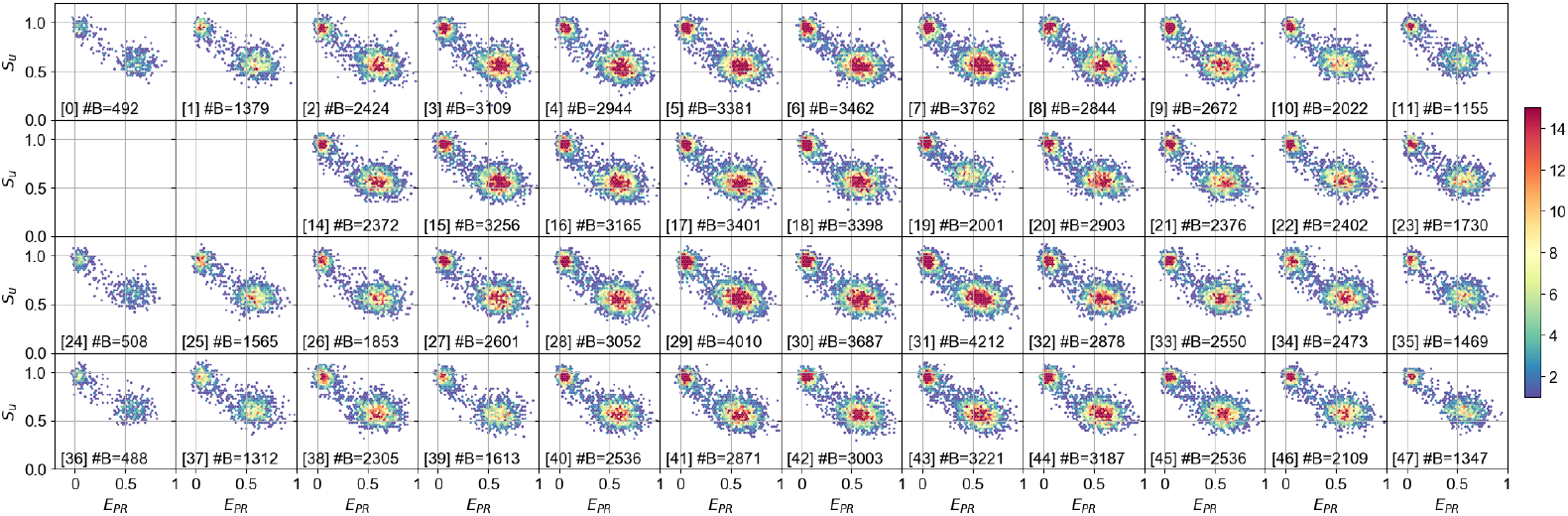
*E_PR_ – S_u_* histograms for each spot from a doubly-labeled dsDNA sample with a 12 bp inter-dye separation. A burst search using all photons was preformed, with m = 10 and a constant threshold of 50 kHz. After background correction, bursts were selected using a minimum burst size of 40 photons. The total number of bursts is indicated as #B at the bottom of each histogram. The 12 bp FRET population (*E_PR_* ≈ 0.6, *S_u_* ≈ 0.6) is isolated from donor-only or acceptor-only populations. For computational details refer to the *48-spot-smFRET-PAX-analysis* notebooks (link). Figure reproduced from ref. [30].

#### 4.1.3. E – S histograms for HT-smFRET-PAX

After burst selection, a 2-D *E* – *S* histogram is plotted, where *E* is the FRET efficiency and *S* is the stoichiometry ratio defined in Appendix C. *S* is approximately equal to *N_D_*/(*N_D_* + *N_A_*), where *N_D_* and *N_A_* are the respective numbers of donor and acceptor molecules present in a burst.

In practice, calculating *E* and *S* exactly requires knowledge of several correction parameters that may not be available at the beginning of the analysis. Instead, related uncorrected quantities that are simpler to compute (*E_PR_* and *S* (or *S_u_*), the latter specific to PAX measurements, defined in Appendix C.6), can be used to identify sub-populations. The corresponding 2-D *E_PR_ – S* or *E_PR_* – *S_u_* histograms allow isolation of FRET species from singly labeled donor-only or acceptor-only species, as shown in Fig. 4.

The accuracy of multispot data analysis was verified by comparing results obtained for each spot. Applying a second burst selection criterion, 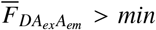 (where the min value is provided in each figure caption) removed the donor-only population and isolated the FRET subpopulations identifiable in Fig. 4. Fitting the corresponding burst distribution with a 2-D Gaussian yields center-of-mass and standard deviation parameters represented in Figure 5A as blue dots and crosses respectively, where the orange dot represents the average position of the FRET peak position over all spots. The overall dispersion of these populations is quite minimal, even without spot-specific corrections, as visible for the FRET population (blue scatterplot in Figure 5B) and the donor-only population (orange scatterplot in Figure 5B).

**Figure 5:**
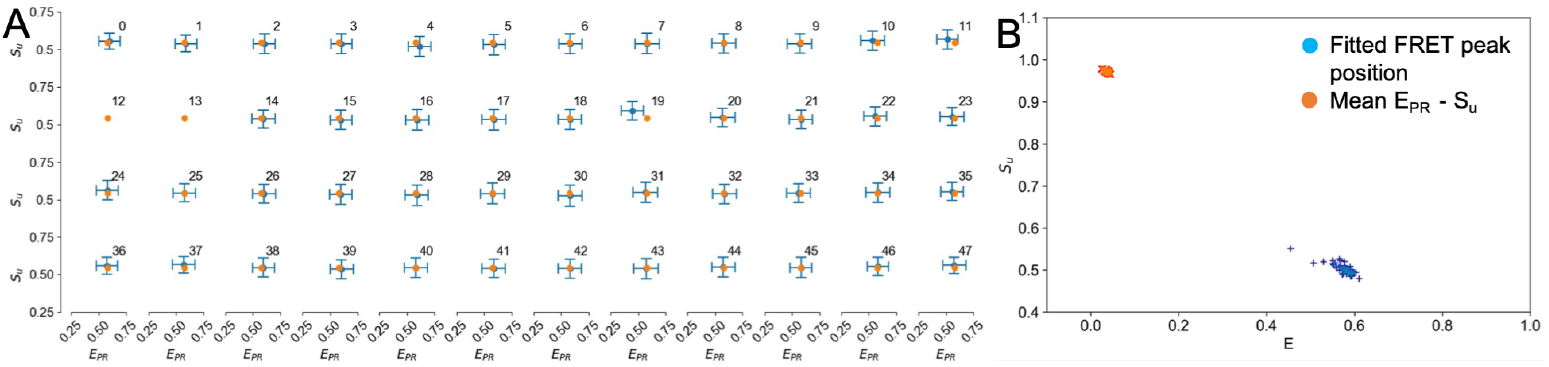
Left: Gaussian fitted *E_PR_ – S_u_* peak position for each spot. FRET peak positions and standard deviations are denoted by blue dots and crosses respectively. Spots 12 and 13 correspond to two defective pixels in the donor SPAD array. Right: FRET (blue crosses) and donor-only (orange crosses) peak position for all spots. For computational details refer to the *48-spot-smFRET-PAX-analysis* notebooks (link). Figure adapted from ref. [30].

#### 4.1.4. Pooling data from HT-smFRET-PAX measurements

The final step of HT-smFRET-PAX analysis involves combining data from each of the spots into a single global data set. Non-uniformities across spots can be accounted for by spot-specific γ and *β* corrections, as discussed in Appendix C.8. Figure 6 shows the result of this process for data obtained from a mixture of doubly-labeled dsDNA with inter-dye distances of 12 bp and 22 bp.

**Figure 6:**
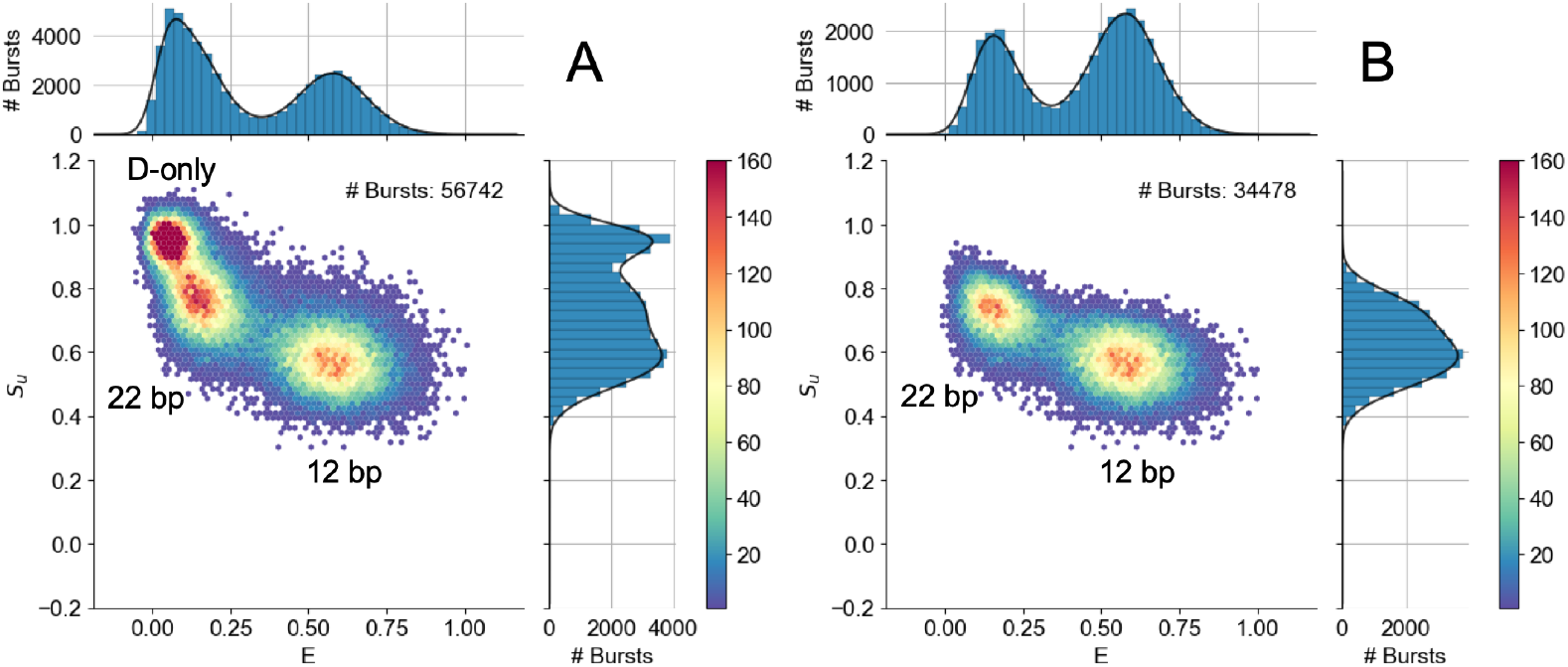
Pooled data from HT-smFRET-PAX measurements of freely diffusing dsDNA separated by 12 bp and 22 bp, where *γ* for the multispot-PAX setup = 0.5. No spot-specific corrections were applied. A) Burst search using *m* = 10 and a constant rate threshold of 50 kHz, burst size selection using > 80 photons. B) Histogram from A) with an additional burst selection, 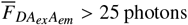. The additional selection removes the donor-only population from the histogram and leaves two FRET subpopulations corresponding to the 12 bp and 22 bp FRET species. For computational details refer to the *48-spot-smFRET-PAX-analysis* notebooks (link). Figure reproduced from ref. [30].

The large number of bursts obtained by this operation allows the use of more stringent selection criteria (e.g. larger minimum burst size) in order to keep only bursts with high signal-to-noise ratio. Additionally, pooling data enables sub-population information to be obtained after a much shorter acquisition time than would be possible with a single-spot setup, as illustrated in Figure 7 where a 5 s acquisition window was used.

**Figure 7:**
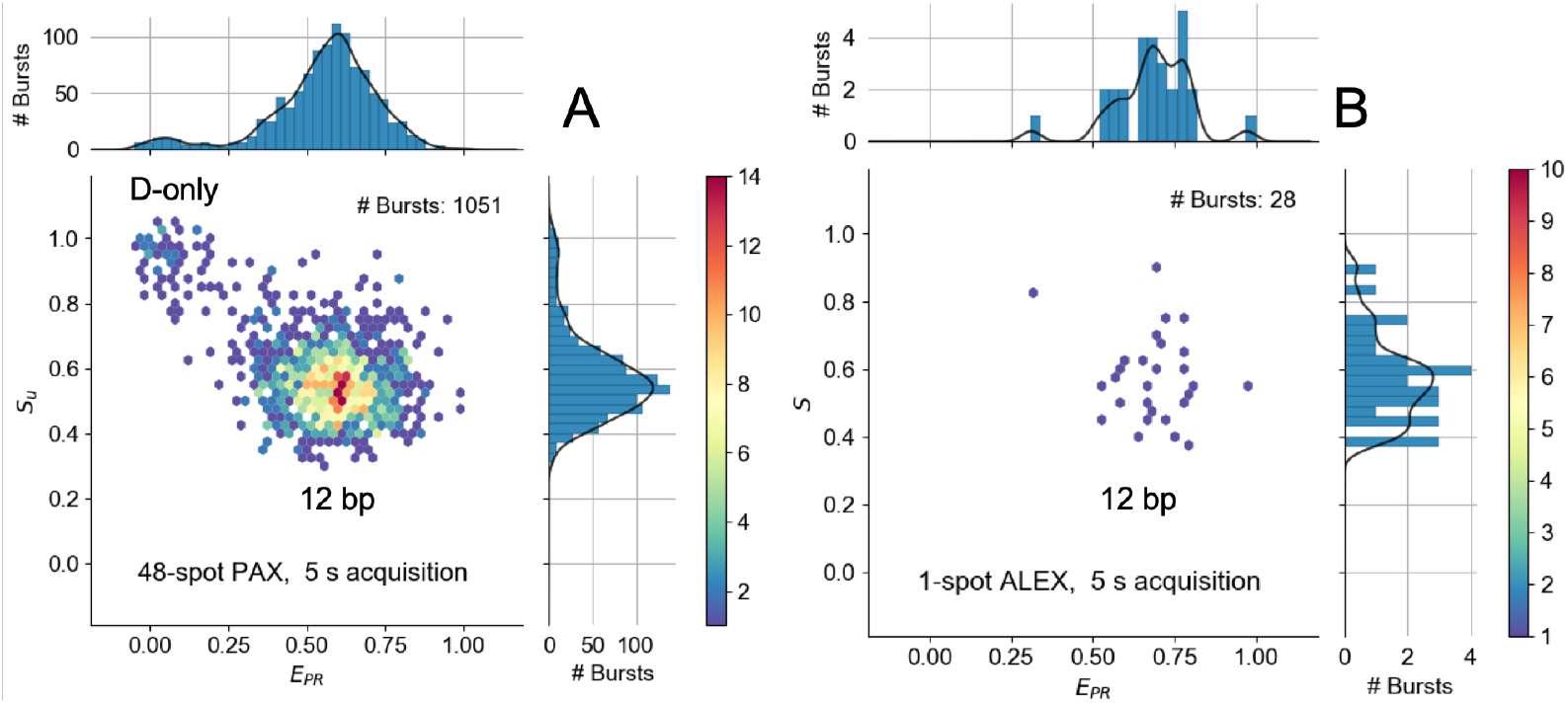
Pooled data from HT-smFRET-PAX measurements of freely diffusing dsDNA separated by 12 bp. Where *γ* for the multispot-PAX setup = 0.5. No per-spot corrections were applied. A) *E_PR_ – S_u_* histogram for 5 s of acquisition with the 48-spot-PAX setup. A constant rate threshold = 20 kHz was applied followed by a burst selection on the counts during donor excitation, 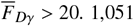 bursts were collected in 5 s. B) *E_PR_ – S* histogram for 5 s of acquisition on the single-spot μsALEX setup using the same sample from A). A constant threshold = 20 kHz was applied followed by a burst selection on the counts during donor excitation, 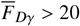. Only 28 bursts were collected in 5 s. For computational details refer to the *48-spot-smFRET-PAX-analysis* notebooks (link). Figure reproduced from ref. [30].

By pooling data from all 46 spots (2 SPADs being defective in one of the arrays), a 38-fold increase in number of bursts is observed in the multispot experiment compared to the single-spot experiment. This ratio fluctuates depending on the observation time point, due to the stochastic nature of single-molecule transit through excitation/detection spots, and to differences between both setups’ excitation/detection volumes and detection efficiencies.

This increased throughput can be used to improve the temporal resolution of out-of-equilibrium reaction studies in “standing drop” sample geometries, where the molecules simply diffuse in and out of the excitation/detection volumes. In theory, the temporal resolution of such a measurement depends inversely on the burst rate (number of bursts detected per unit time). However, this is only true long after the reaction is well established throughout the sampled volume, as will be discussed in the next section.

### 4.2. Kinetic study of bacterial transcription

We used our original 8-spot setup to study the kinetics of transcription initiation by bacterial RNA polymerase (RNAP) as a simple demonstration of high throughput multispot smFRET [29], as described next.

#### 4.2.1. Bacterial transcription initiation

DNA transcription into RNA by RNAP occurs in three main steps: initiation, elongation, and termination. Transcription initiation is highly regulated and is the rate limiting step of the reaction [65]. This stage is comprised of four steps, involving:

1. binding of the core RNAP by the promoter specificity σ factor to form the RNAP holoen-zyme,
2. binding of RNAP holoenzyme to DNA at the promoter sequence upstream from the gene sequence to form the RNAP-promoter closed bubble (*RP_c_*) complex,
3. a multistep sequence of events leading to the unwinding of 10-12 bp of the dsDNA promoter sequence and formation of the RNAP-promoter open bubble (*RP_o_*) complex, and finally,
4. the initial polymerization of RNA, involving many failed attempts (abortive initiation), ultimately leading to promoter escape and the transition to elongation.

The open-bubble can be stabilized by the addition of a dinucleotide, corresponding to the so-called *RP*_*ITC*=2_ intermediate complex (Figure 8A). The *RP*_*ITC*=2_ complex is stable until nucleotides (nucleoside triphosphates, NTPs), necessary to transcribe the gene, are added [66].

**Figure 8:**
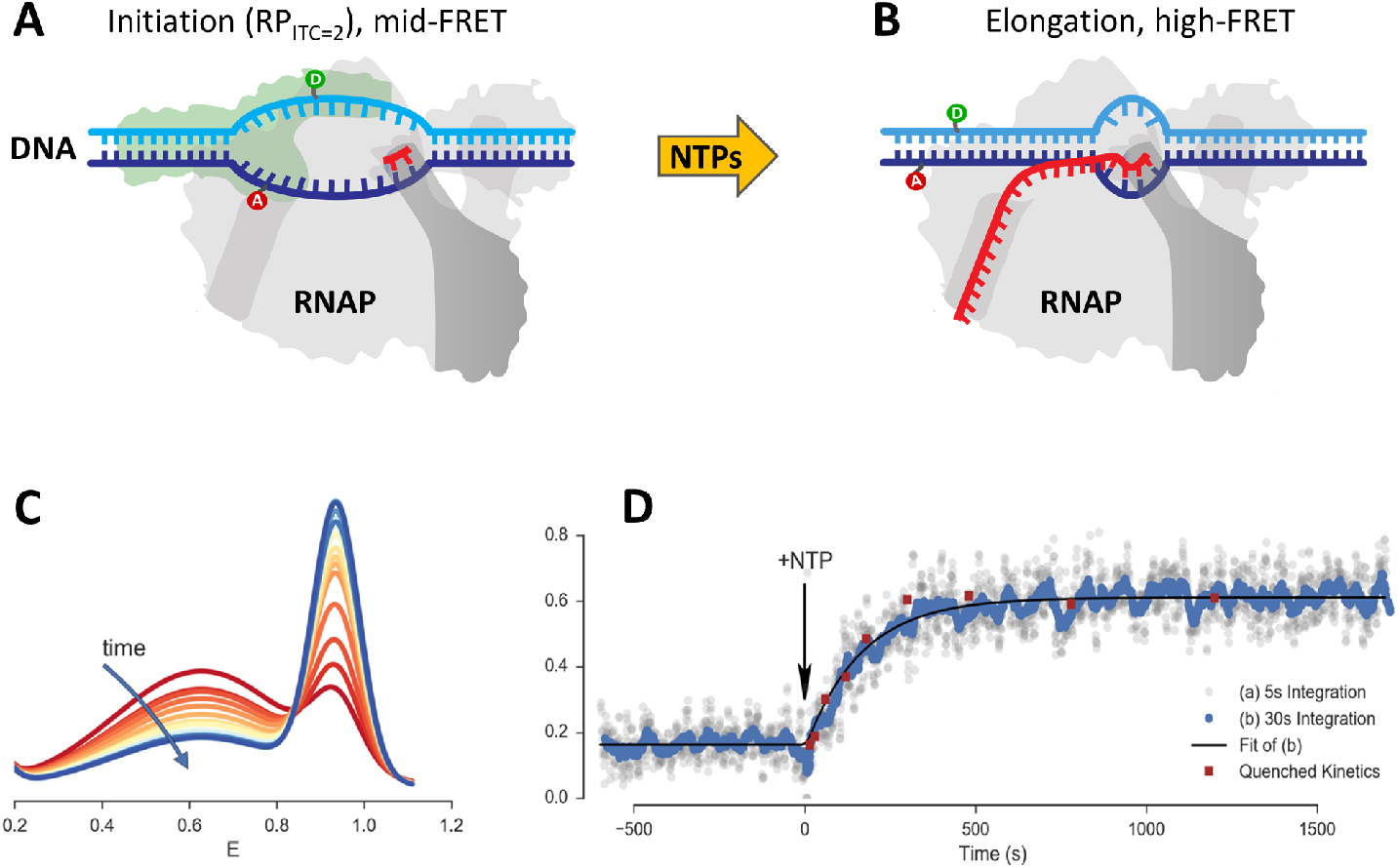
RNAP kinetics study. A) The RNAP-promoter initial transcribing complex *(RP*_ITC_) is prepared with a stabilizing dinucleotide (red symbol) as the nascent RNA chain. Complementary DNA strands are labeled at DNA promoter bases with donor (D, green, position −5) and acceptor (A, red, position −8) dyes. After formation of a transcription initiation bubble, the bases to which the dyes are conjugated are separated, resulting in medium FRET. The initial state remains in stationary conditions until the addition of the four missing nucleotides (NTPs, yellow arrow), which triggers transcription initiation and elongation. B) During elongation, the transcription bubble moves downstream (to the right), resulting in re-hybridization of the open-bubble sequence at the promoter sequence, and a corresponding decrease of the D-A distance (*i.e.* a FRET increases). C) Evolution of uncorrected FRET efficiency (*E_PR_*) distributions as function of time. The curves represent Gaussian fits of the *E_PR_* histograms using 30 s time windows. D) Fraction of high FRET population obtained in the real-time kinetics measurement (grey and blue dots). Dots are computed as a function of time using either a 5 s (grey) or 30 s (blue) moving integration window. The solid black curve is a single-exponential model fitted to the 30 s moving integration window. Quenched kinetics data (red dots) [66], normalized to fit initial and final values of the real time kinetics trajectory, are also shown for comparison. For more details on the analysis see the Jupyter notebook provided in ref. [29]. Figure adapted from ref. [29].

The transcription reaction begins after all four NTPs (ATP, UTP, GTP, and CTP) are added to the assay. Even after formation of the *RP_o_* complex, further initiation steps can postpone the transition to elongation, and hence are rate limiting to the transcription process. In abortive initiation, short transcripts are formed with the assistance of RNAP scrunching DNA into the bubble available to the transcription active site. However, due to sigma region 3.2 blocking the RNA exit channel, nascent transcripts are backtracked with the assistance of DNA un-scrunching, until the short RNA is released. Ultimately, this leads to multiple failed transcription attempts. Additionally, transcription pausing can further postpone the transition from transcription initiation to elongation, as elucidated by smFRET and single-molecule magnetic tweezer experiments [67, 68]. It is only when the blockage of the RNA exit channel is relieved that the transition from initiation to elongation proceeds, followed later on by termination.

Termination is characterized by a closed bubble where the *RP_o_* complex was initially located (Figure 8B). To characterize this transition, we studied its kinetics from the *RP*_*ITC*=2_ stage until promoter escape occurred. FRET between two labeled nucleotides on opposite strands in the bubble region can be monitored, e.g. between the template strand labeled with ATTO 550 (donor) and the non-template strand labeled with ATTO 647N (acceptor). In *RP*_*ITC*=2_ (initiation stage), the dyes are separated (medium FRET), while during or after elongation, the DNA strands at the promoter sequence re-anneal, leading to a small inter-dye distance characteristic of dsDNA, and therefore high FRET.

#### 4.2.2. RNAP kinetics

We monitored the initial stage of transcription using the 8-spot HT-smFRET setup described above [29] by triggering the reaction with manual addition of a full set of nucleotides, preceded and followed by continuous recording of smFRET bursts from diffusing single complexes in solution (experimental details can be found in ref. [29]).

Data analysis was performed essentially as for a steady state or equilibrium measurement, using standard background estimation, burst search and burst selection procedures (but no corrections), with the only difference that the resulting bursts where grouped in different windows as described next. The initial part of the experiment prior to nucleotide addition was used to identify the two sub-populations (*RP*_*ITC*=2_: medium FRET, *E_PR_* = 0.62, and RNAP-free DNA: high FRET, *E_PR_* = 0.95), as a fraction of free DNA population is expected in these measurements. This free DNA population is indeed indistinguishable from the final population of molecules having undergone complete transcription. After nucleotide addition, the burst population was analyzed in 30 s windows moved with 1 s increments, and the resulting FRET sub-populations were characterized by their fractional occupancy as a function of time (Figure 8C).

A clear first order exponential kinetics is observed in Figure 8D, characterized by a lifetime *τ* = 172 ± 17s. This behavior matches that observed using a completely orthogonal approach involving a series of quenched transcription reactions monitored by standard equilibrium smFRET measurement in solution [66] (red dots in Figure 8D), validating this HT-smFRET approach for slow kinetics. Interestingly, data analyzed with 5 s sliding windows (grey dots in Figure 8D) exhibit the same trend, although with a smaller signal-to-noise ratio, confirming the importance of as large a number of sampling volumes as possible in order to access short time scales. However, the resolution of this relatively crude approach of triggering the reaction based on manual addition and mixing of reactants is limited by the dead-time of the mixing process itself, on the order of 10-20 s in these measurements. Accessing shorter time scales will require combining this approach with automated and faster microfluidic mixing.

### 4.3. HT-smFRET in microfluidic devices

#### 4.3.1. Introduction

As suggested above, multispot SPAD arrays will find their full potential in high-throughput applications when combined with microfluidic devices. In the following, we present three types of devices that enable different types of HT-smFRET:

1. Microfluidic “formulator” device,
2. Parallelized microfluidic device,
3. Microfluidic device based on hydrodynamic focusing.

The microfluidic “formulator” device (Fig. 9A) [69, 70] allows rapid mixing of reactants with picoliter (*pL*) precision, measurement for an extended period of time, sample flushing, and automated titration for an arbitrary number of repetitions. HT-smFRET analysis in such a device would extend the throughput of previous measurements limited to single-spot geometry [70], and allow rapid study of the equilibrium conformational landscape of biomolecules or mapping of the dependence of enzymatic activity as a function of its chemical environment.

**Figure 9:**
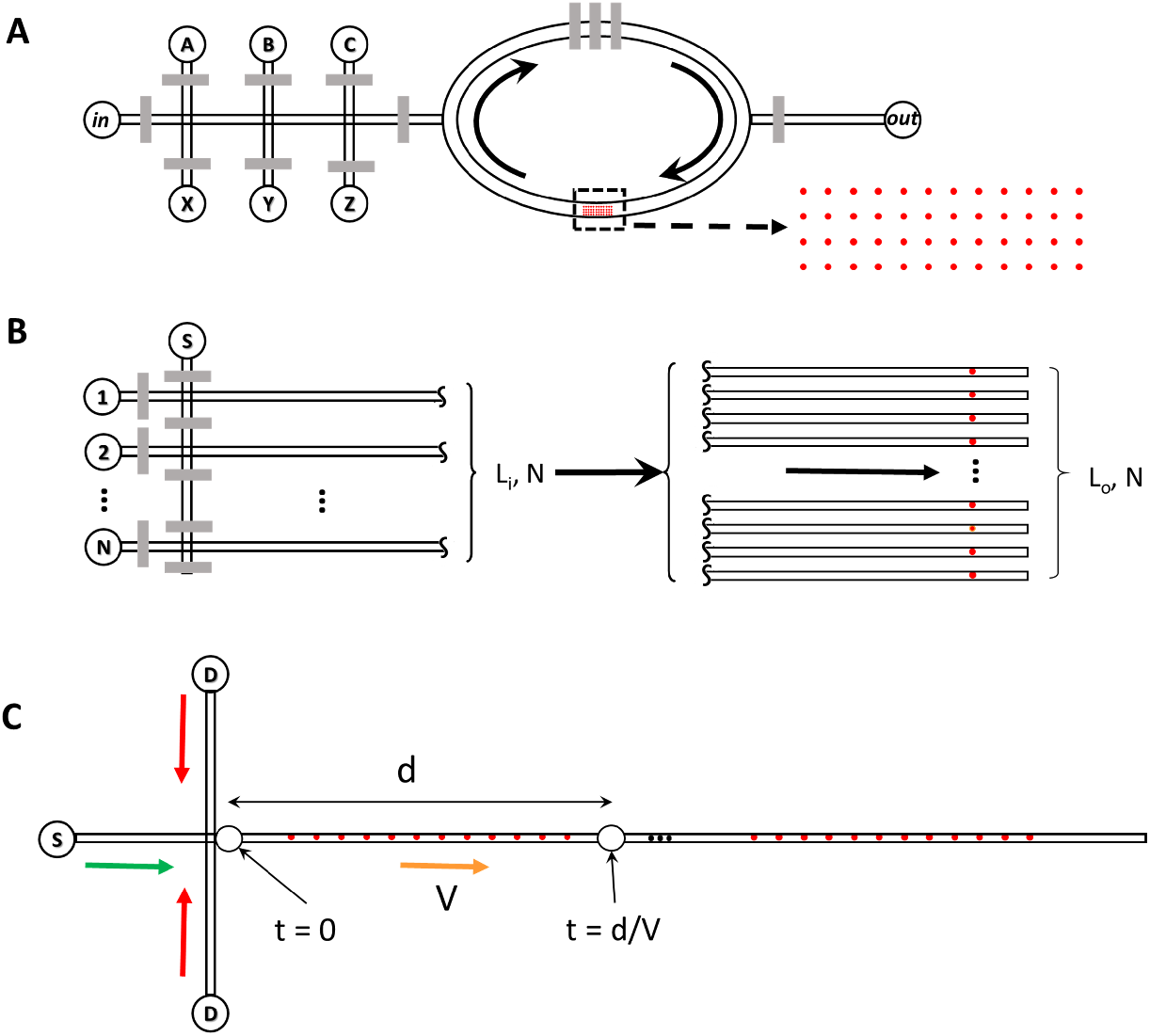
Examples of possible SPAD array and microfluidic combinations A) Formulator geometry: in a microfluidic formulator, several sample reservoirs (A, B, C, …, X, Y, Z) are connected to an injection channel via programmable valves (grey rectangles), which allow precise injection of volumes (*pL*) of sample within a mixing region (ellipse). A peristaltic pump mechanism (top three grey rectangles) allows mixing the different sample aliquots within a few seconds. Equilibrium measurements of the mixed sample can then be performed in an observation chamber (dashed rectangle) using a dense array of independent spots such as that described in section 4.1.1. B) High-throughput screening geometry: a linear multispot array combined with a multichannel microfluidic channel would allow high-throughput, parallel singlemolecule detection, with clear applications in molecular screening and diagnostics. The channel separation on the inlet side (*L_i_/N*) is much larger than their separation on the outlet side (*L_o_/N*), set to match the excitation spot pitch. C) Microfluidic mixer geometry: in a fast single-molecule microfluidic mixer, a sample (S) is rapidly mixed with another solution (D), plunging molecules quasi-instantaneously in a different environment, thus triggering a series of changes (conformation, chemical, or enzymatic reaction, etc.). A multispot setup with a linear illumination scheme and SPAD array would allow acquiring information from individual molecules at as many time points along the reaction coordinate in parallel, thus speeding up fast kinetic measurements.

In contrast to the experiments presented in ref. [70] where the (single-spot smFRET) measurement time was the limiting factor (resulting in overnight data acquisition duration), HT-smFRET could bring the measurement time down to the mixing time scale of this type of mixer (a few s). For example, if one data point is collected by an 8-spot setup in a 5 s acquisition time window (as described in Fig. 8), using our 48-spot setup would bring the required acquisition time to achieve similar statistics down to less than 1 s. This time resolution is much faster than what can be achieved with a standard single-spot setup, but still much slower than what can be achieved with a continuous flow mixer, which can reach millisecond time scales, as discussed in Section 4.3. In addition to speeding up acquisition and therefore making this approach a practical analytical tool rather than just a research tool, reduced experiment duration would have several other advantages, such as reduced sample degradation and setup drift.

A parallelized microfluidic approach is implemented in Fig. 9B, in which each spot of a multispot setup probes a unique sample. This device could be comprised of many independently addressable channels with the use of a microfluidic multiplexer, allowing, for instance, probing a common sample (S) with multiple probes (1… N) after controlled mixing. This parallel geometry is more technically challenging because it requires a good match between spot density (limited by the field of view of a high numerical aperture microscope) and microchannel density (limited by the resolution of soft-lithography). This approach may require custom-designed optics for larger SPAD arrays than those described in this article.

Microfluidic hydrodynamic focusing (Fig 9C) [71, 4, 72, 73] achieves mixing rates orders of magnitude faster than the formulator design described above, by injecting a sample (S) into a cross-junction carrying a “diluent” solution (D) such that the three input streams are mixed in the outlet channel (other geometries accomplishing the same goal are also possible, for example see ref. [5]). As long as the flow remains laminar, the net result is a thin (<< 1 *μm*) slab of sample S focused between laminar streams of surrounding diluent solution. Due to the small width of the sample slab, sample and solute molecules diffuse and mix on the timescale of microseconds to milliseconds (*μs* – *ms*). Past this “time 0” point within the mixer’s main channel, sample molecules evolve in a diluent environment as they flow along the main channel, the time *t* since the start of the reaction being given by *t* = *d/V*, where *d* is the distance from the mixing region and *V* the is the flow velocity. Single-molecule measurements with hydrodynamic focusing typically used single-spot approaches and require accumulation of data one time-point at a time, which is both time and sample consuming. A linear SPAD array geometry, combined with a linear illumination pattern such as demonstrated in ref. [17] would significantly speed up data acquisition in this type of fast kinetics experiment, as well as offer the exciting possibility of tracking the evolution of individual molecules along their reaction path.

Similar kinds of measurements have been previously demonstrated using cameras and have achieved temporal resolution on the order of 100 *μs* [74, 75] to 10 *ms* [76]. In these geometries, single-molecules are either flown rapidly (> 50 *mm/s*) in a simple microfluidic channel and detected as streaks by an electron-bombarded camera [74, 75], or flown in very narrow channels at speeds compatible with single-molecule localization (< 1 *mm/s*) and tracked by stroboscopic illumination [76].

Both approaches could be used with SPAD arrays and fast single-molecule microfluidic mixers. In the first case, fast flow (> 50 *mm/s*) would result in very low counts per spot, due to the limited excitation intensity and short transit time, but high likelihood to detect the same molecule in consecutive spots along the flow direction, due to the large flow velocity minimizing the effect of lateral diffusion. With a spot separation of 2 *μm*, a 40 *μs* resolution would be obtained. Such a resolution would approach that of ultrafast mixer measurements using non-single-molecule concentrations [71, 77, 72]. While these individual single-molecule time traces would provide little information due to the low signal to noise ratio (SNR), cumulative statistical analysis of large numbers of such time traces could provide unprecedented insights into single-molecule dynamics.

In the second case (flow velocity < 1 *mm/s*), time resolution of identical spot separation would be reduced to > 2 *ms*, but the SNR would remain compatible with single time point FRET measurement. However, the likelihood of capturing the same molecule in consecutive spots would be limited by diffusion. Nonetheless, cumulative analysis of large numbers of such short trajectories would likewise provide unique information on single-molecule dynamics.

#### 4.3.2. Proof of principle experiment

A simple microfluidic device with a single channel containing a viewing chamber of dimensions *L* × *W* × *H* = 3.6 *mm* × 320 *μm* × 10 *μm* mounted on a glass coverslip was used to test the compatibility of multispot HT-smFRET with flow. Inlet and outlet holes (≈ 0.5 *mm* in diameter) were created using a biopsy punch, and connected to 20 gauge Tygon tubing by 23 gauge stainless steel pins. The outlet tubing was connected to a luer-locking 23 gauge syringe tip connected to a 1 mL Norm-ject syringe mounted in a programmable syringe pump (NE-1000 Multi-Phaser, New Era Pump Systems, NY). A 500 pM sample of doubly-labeled dsDNA sample (ATTO 550 and ATTO 647N separated by 5 bp [78]) was injected into the inlet Tygon tubing and pulled into the chip with the syringe pump at a constant flow rate of ~ 10 *μL/hr*. The microfluidic chip was installed on the 48-spot smFRET-PAX setup discussed in section 4.1, and measurements were performed using a total laser power of 500 mW for the 532 nm and 800 mW for the 628 nm laser in the presence of flow (average power at 628 nm due to alternation: 400 mW). Control experiments performed on a standing drop used lower powers (532 nm: 300 mW, 628 nm: total 600 mW, average 300 mW) to account for the shorter residence time of molecules in the excitation spots in the case of flow.

#### 4.3.3. Flow characterization by CCF analysis

Flow velocity can be extracted by computing the CCF of the intensity signals recorded at two locations separated by a distance *d* along the flow direction (two-beam cross-correlation) [79]. The normalized 2D CCF takes the form:

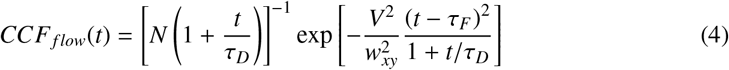

where *τ_D_* is the diffusion time across each excitation/detection volume, assumed Gaussian in *x* – *y* with waist *w_xy_* (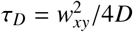, where *D* is the diffusion constant), *V* is the flow velocity and *τ_F_* = *d/V* is the time it takes a molecule to traverse the distance between two adjacent spots.

In the geometry of this measurement (Figure 10A), there are 36 pairs of spots separated by *d*_0_ = 5.4 *μm*, 24 pairs of spots separated by 2 × *d*_0_ and 12 pairs of spots separated by 3 × *d*_0_ (pairs at an angle with respect to the flow direction could also be considered for this analysis). Since they are equivalent, it is possible to average CCF’s corresponding to the same separation but different pairs, resulting in the curves shown in Figure 10B. In the presence of flow, peaks at characteristic time scales *τ_F_i__* (*i* = 1,2,3) ~ 21,41,60 *ms* are visible in both channels along the direction of the flow, but not in the opposite direction, as expected. By comparison, no peak is detected in the absence of flow (Figure 10C). The translation time between consecutive spots corresponds to an average flow velocity *V_meas_* ~ 257 *μm/s*), slightly different from that corresponding to the programmed flow rate and channel dimension (*V_theo_* ~ 309 *μm/s*), but consistent with that expected at a slightly off-center vertical position due to the quasi-parabolic dependence of the velocity profile with the vertical position within the channel [80].

**Figure 10:**
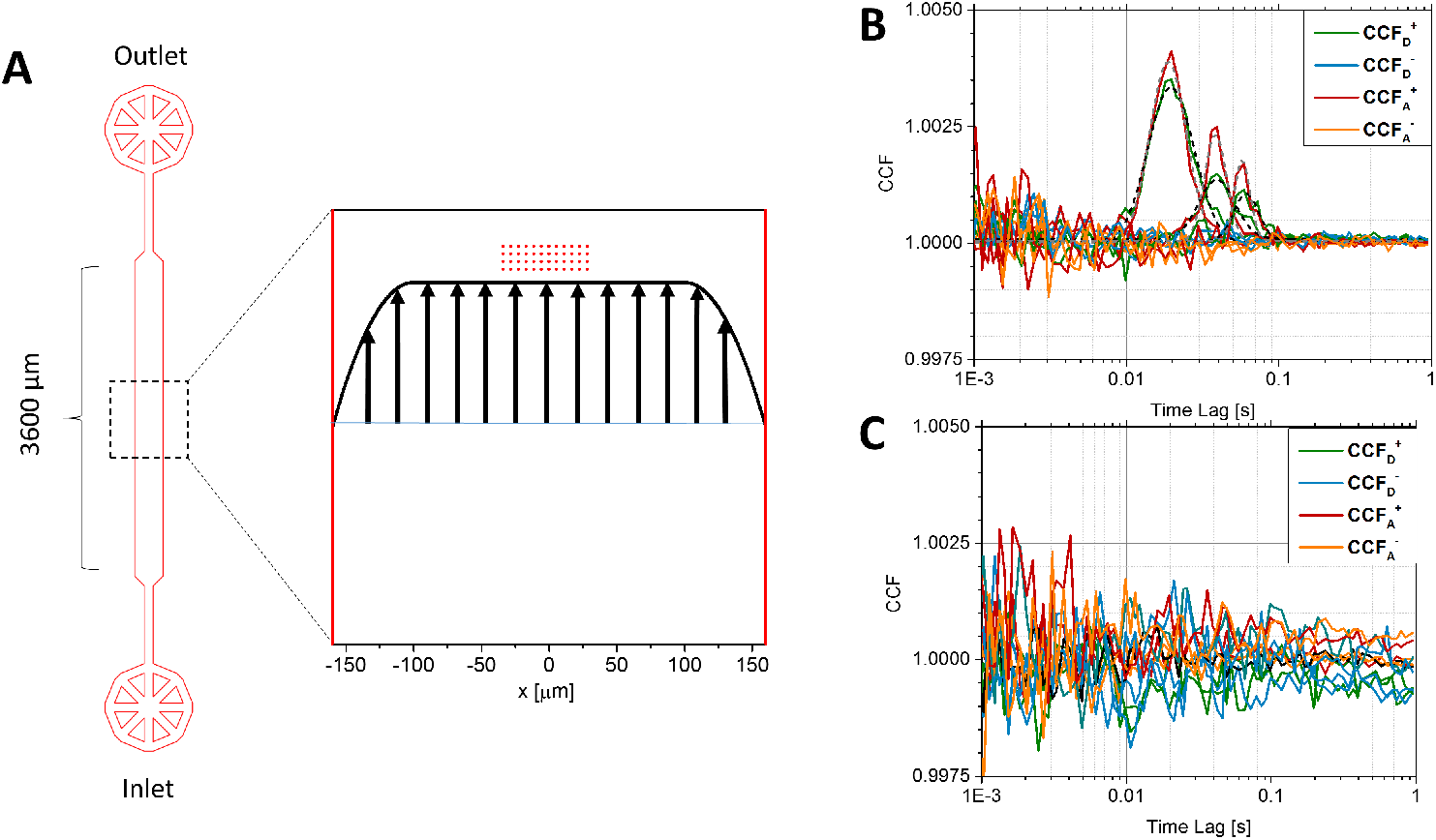
HT-smFRET in a microfluidic chip. A: Measurements were performed in the center of a simple microfluidic chamber in which a single-molecule sample of doubly-labeled dsDNA lacCONS promoter sequence (ATTO 550 at the −3 register with respect to the transcription start site and ATTO 647N at −8 bp from the start site [78]) was flown at a constant flow velocity *V*. The 48-spot excitation pattern (red dots, width ~ 60 *μm*, pitch distance *d*_0_ = 5.4 *μm*) was located in the center of, and perpendicularly to the 320 *μm*-wide channel, in a region where the velocity profile (schematically represented by the black curve and parallel arrows) is approximately constant in the *x – y* plane and parabolic along the vertical direction (not shown). B: The average CCFs of adjacent spots (distance *d*_1_ = *d*_0_), spots separated by *d*_2_ = 2 × *d*_0_ or *d*_3_ = 3 × *d*_0_ along the direction of the flow, calculated for both donor (green) and acceptor (red) detection channels over the first 200 s, exhibit a clear peak around *τ_F_i__* ∼ 21,41 and 61 *ms*, as fitted using Eq. 4 (grey and black dashed curves). No such peak is visible in the corresponding average CCFs computed in the reverse direction (blue and orange curves) or in the absence of flow. Fits of the CCF curves with Eq. 4 (dashed curves) yield an average flow velocity *V* = 253 ± 6 *μm/s*, or a transit time across a single spot *τ_T_* ~ 3*ω_x_y*/*V* ∼ 3 *ms* ~ 11*τ_D_*, where the diffusion time *τ_D_* ∼ 268 *μs* was obtained from a fit of the average donor channel ACFs. Datasets used for this figure as well as ALiX notebooks and associated files used for analysis can be found in the Figshare repository [81].

#### 4.3.4. HT-smFRET in a simple microfluidic device

The measured velocity is within the range of flow velocities used for smFRET analysis in microfluidic mixers [4, 73], which requires a transit time long enough to accumulate a sufficient number of photons during a single-molecule burst. It is however much smaller than velocities used for high-throughput single-molecule detection (several cm/s), which require much higher excitation powers to obtain a detectable single-molecule signal [82].

To assess the effect of flow on single-molecule burst characteristics, we compared the *E_PR_ < S* histograms, pooled over the 48 spots, obtained first for the sample observed in conditions of free diffusion, and next in the presence of flow (but excited with higher power, see above), recorded over a common duration of 200 s (Figure 11). While the relative fractions of donor-only and FRET bursts is different due in part to the different excitation intensities used in both measurements, their *E_PR_* and *S* characteristics are identical.

**Figure 11:**
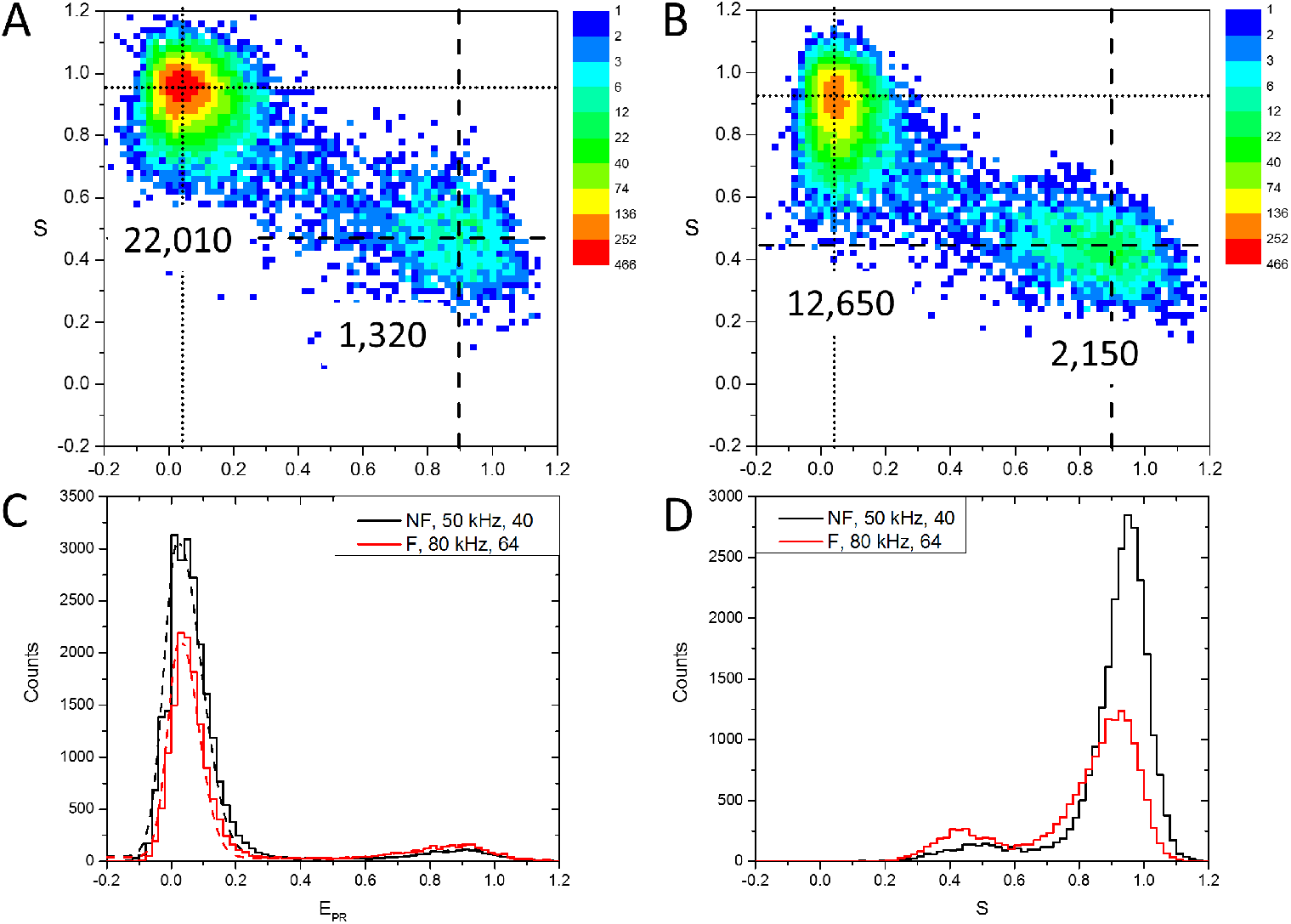
A,B: Comparison of the *E_PR_ – S_u_* histograms of a dsDNA FRET sample in the absence (A) or in the presence (B) of flow. The diffusion only (no flow or NF) dataset, recorded with lower excitation powers (by a factor ~ 1.6), was analyzed with a lower rate threshold (*r_m_* ≥ 50 *kHz*) for burst search and a lower burst size threshold 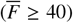 for burst selection, than the dataset recorded with flow (F), for which *r_m_* ≥ 80 *kHz* = 50 × 1.6, 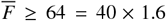, in order to obtain comparable number of bursts for analysis. The numbers next to each sub-population (top-left: donor-only, middle-right: FRET) correspond to the estimated integral under each peak as discussed in C. The *E_PR_, S_u_* location of the donor-only and FRET populations is identical in both experiments. Note that the color scale is logarithmic. C: Projected *E_PR_* histograms for the no flow (NF, black) and flow (F, red) measurements. Dashed curves correspond to fits with a model of asymmetric Gaussian with tilted bridge described by Eq. 19 in section C.7. The integral under each peak, given by Eq. 20 provides an estimate of the number of bursts in each sub-population, as reported in A & B. D: Projected *S_u_* histograms for the no flow (NF, black) and flow (F, red) measurements. Details of the analysis can be found in the different notebooks: `ALiX Notebook, XX, APBS, m = 10, Rmin = YY kHz, Smin = ZZ.rtf` where XX = Flow or No Flow, YY = 50 or 80, ZZ = 40 or 64, and associated files in the Figshare repository [81]).

The effect of using different powers can be partly mitigated by using a burst search rate criterion (*r_m_* > *R_min_*, see Eq. 14) and burst selection criterion 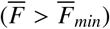, adjusted proportionally to the excitation power. The donor laser excitation power is for instance reflected in the burst peak count rates of the D-excitation, D-emission photon stream (Figure 18).

This increase implies that the throughput (number of bursts recorded per unit time) of measurements in equilibrium conditions can be greatly increased even by modest flow rates, a concept already demonstrated in single-spot geometry [82]. Moreover, contrary to diffusion-only measurements, each burst observed in a given spot in the presence of flow corresponds to a different molecule, rather than potentially to the same recurring molecule diffusing in and out of that spot. The resulting statistics can thus be directly translated into true sample concentration characteristics, without the uncertainty due the stochastic nature of the number of bursts per molecule detected in diffusion-only experiments.

Analysis of other statistics, such as burst size or burst duration is complicated by the different excitation power used in both measurements. However, the burst peak count rates of the donor-only and acceptor-only populations can be compared: they indeed scale as the excitation powers used for each experiment. These results clearly indicate the potential of combining HT-smFRET and microfluidics, although a number of trade-offs will need to be studied in future work. For instance, while burst numbers would first increase at higher flow velocity, the shorter translational transit time (*τ_D_*) would eventually be accompanied by lower burst peak count rate, which, unless compensated by different burst search and selection criteria, and by increased excitation power, would eventually result in decreasing numbers of detected bursts [82]. Moreover, increased excitation will result in increased photobleaching [83, 84, 85], especially in mixer geometries, where the same molecule may cross several spots successively (and be excited continuously for a long period of time in the case of linear illumination geometry). Being able to follow the evolution of the same single molecule across successive spots would however open up fascinating perspectives to study fast conformational dynamic trajectories.

## 5. Conclusion and perspectives

Over the past decade, the development of SPAD arrays with performance compatible with smFRET measurements has opened up a number of exciting possibilities for high-throughput single-molecule fluorescence measurements. While there is still room for improvement in terms of detector sensitivity (partially achieved with red-enhanced SPAD arrays) and lower dark count rate, the characteristics of current arrays (both in terms of sensitivity and number of SPADs) already allow envisioning several extension of this work into equilibrium HT-smFRET measurements using sophisticated microfluidic formulator devices, HT-smFRET kinetics using fast microfluidic mixers and high-throughput screening using parallel channel microfluidic lab-on-chip devices (Fig. 9) [86, 87].

This combination will probably require specialized microfluidic designs to take advantage of, and accommodate the new SPAD arrays, and in turn, motivate new SPAD array geometries for specific applications. In particular, fast microfluidic mixer or parallel channel high-throughput screening applications would benefit from linear SPAD arrays with larger number of SPADs and higher density.

Extension of this type of measurements to time-resolved detection is not only possible, as shown above, but most desirable, as it provides information on fast interconverting subpopulations, which are key to understanding dynamic phenomena occurring on time scales shorter than the typical diffusion time, as well as facilitating the detection of short transient states [3].

On the optics side, multispot excitation approaches using spatial light modulators, as illustrated in this work, could potentially be replaced by simpler and cheaper illumination schemes such as the linear illumination approach used in ref. [17]. This would not only facilitate alignment and wider adoption, but also allow more efficient use of laser power, thus lowering excitation power requirements (and cost).

Twenty years after the first demonstration of smFRET measurement in solution [2], there is still a lot to expect from this powerful technique indeed [3].

## 6. Acknowledgments

We are grateful for the contribution of former lab members to the early developments of this technology as well as that of early users of the HT-smFRET setup. We also gratefully acknowledge the work of our POLIMI collaborators whose detectors have made this work possible and will fuel further progresses in years to come. We thank Mrs. Maya Lerner for preparation of illustrations for Fig. 8, panels A & B.

This work was supported in part by NIH grants R01 GM095904, R01 GM069709, R01GM130942, by NSF awards MCB 1244175, MCB 1818147, EAGER 20190026, and by a seed grant from the UCLA Jonsson Comprehensive Cancer Center. S. Weiss discloses intellectual property used in the research reported here.

## Appendix A Data and software availability

### A.1 Data

Previously published raw data, scripts and other files used in this article can be found in free online repositories referred to by their DOI in each figure. New datasets and analysis files specific to this article can be found in a dedicated Figshare repository [81].

### A.2 software and analysis results

Software used to analyze the data presented in this article is freely available online at links provided in the caption of each figure or in the main text and appendices. Specifically, FRETBursts can be found at https://fretbursts.readthedocs.io/en/latest/ and ALiX at https://sites.google.com/a/g.ucla.edu/alix/. The free OriginViewer software (https://www.originlab.com/viewer/ can be used to view the Origin project file (.opj) containing the plots and result of fits shown in Fig. 10, 11, 14, 15, 16, 17, 18 and 19. In addition, the FRETBurst analysis notebooks are available on Github (link).

## Appendix B Setup description and parts information

The make and model of the parts used in the 48-spot setup are provided here for researchers interested in building a multi-excitation wavelength, multispot microscope based on LCOS-SLMs. While at the time of this writing, no single-molecule sensitive SPAD array is commercially available, we hope that this will become the case in the future.

A schematic of the 48-spot setup is presented in Figure 12. The setup uses two 1 W CW lasers (2RU-VFL-Series, MPB Communications, Inc., QC, Canada) with excitation wavelengths 532 nm (green) and 628 nm (red). Both laser intensities can be controlled by software or using a polarizer.

**Figure 12:**
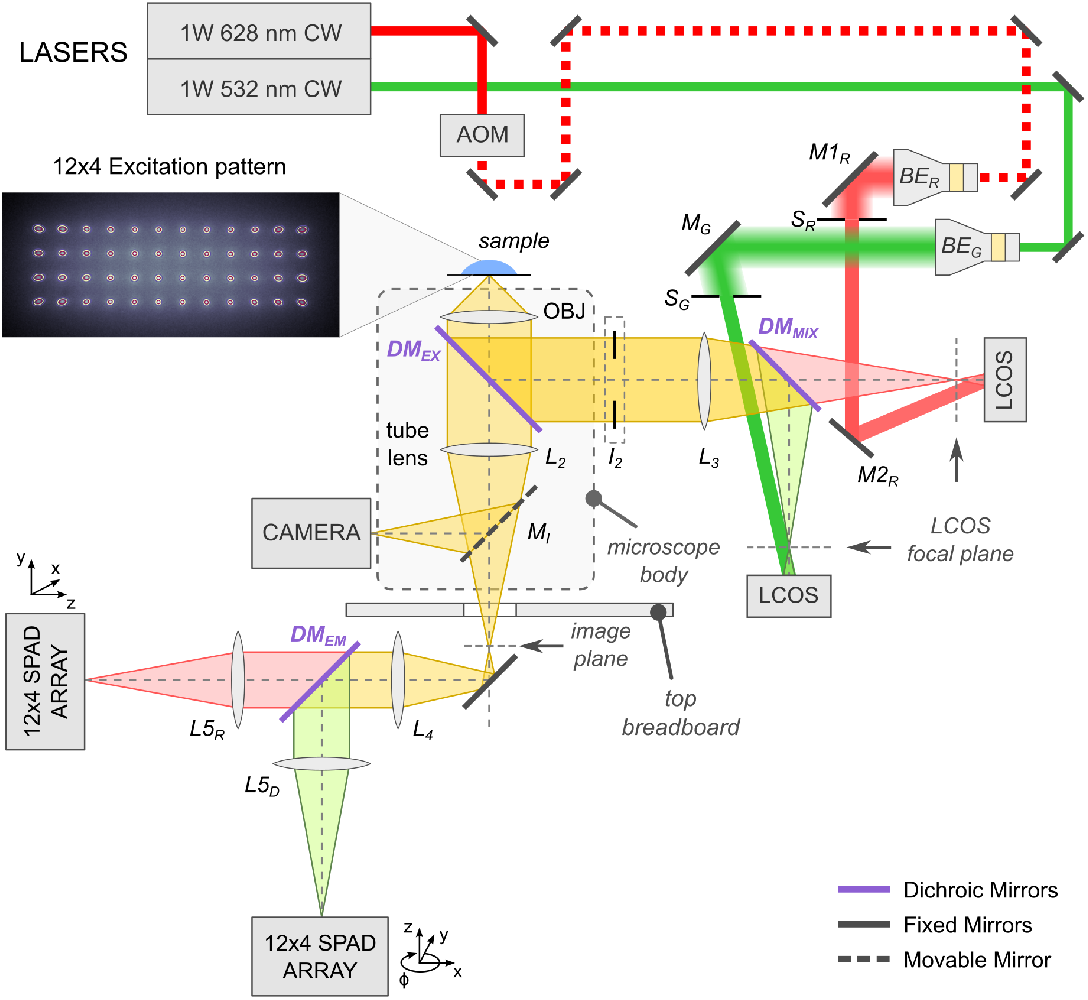
48-spot PAX design schematic. The two CW lasers and SPAD arrays are fixed to a floated optical table. Periscopes are used to bring the beams to the optical breadboard supporting the microscope and LCOS-SMLs. Two beam expanders, mirrors, one dichroic mirror and one lens are used to steer the beams to their respective SLMs, form spot arrays and relay them to the back of the back of the microscope objective lens. The microscope side port is used to monitor the beam pattern using a CMOS camera (an example of which is shown on the left), while the bottom port is used to send the fluorescence signal to the two SPAD arrays via relay lenses, a dichroic mirror and emission filters. A detailed description can be found in the text. Reproduced from ref. [30].

The red laser is alternated by an acousto-optic modulator (AOM; P/N 48058 PCAOM, electronics: P/N 64048-80-.1-4CH-5M, Neos Technology, Melbourne, FL) driven by a square wave (TTL) with 51.2 *μs* period (50 % duty cycle). The polarization of both lasers is controlled by a separate half-wave plate, to match the expected polarization at the SLMs.

Both laser beams are first expanded and collimated using a pair of doublet lenses (Keplerian telescope, with focal lengths *f*_1_ = 50 mm and *f*_2_ = 250 mm, not shown). The laser beams are then steered up to the optical breadboard supporting the microscope using two periscopes and further expanded using two adjustable beam expanders (*BE_G_* and *BE_R_*: 3X, P/N 59-131, Edmund Optics).

Each expanded beam is then steered with mirrors *M*1_*R*_ and *M*2_*R*_, respectively, toward its respective SLM (green: P/N X10468-01, Hamamatsu, Japan, red: P/N X10468-07), which forms an array of spots at its focal plane (Figure 13). Light emitted from these spots is first combined with a dichroic mirror, *DM_mix_* (T550LPXR, Chroma Technology, VT) and focused on the microscope object plane using a collimating lens *L*_3_ (*f* = 250 mm, AC508-250-A Thorlabs) and a water immersion objective lens (UAPOPlan NA 1.2, 60X, Olympus). A dual band dichroic mirror, *DM_EX_* (Brightline FF545/650-Di01, Semrock, NY), is used to separate excitation and emission light.

**Figure 13:**
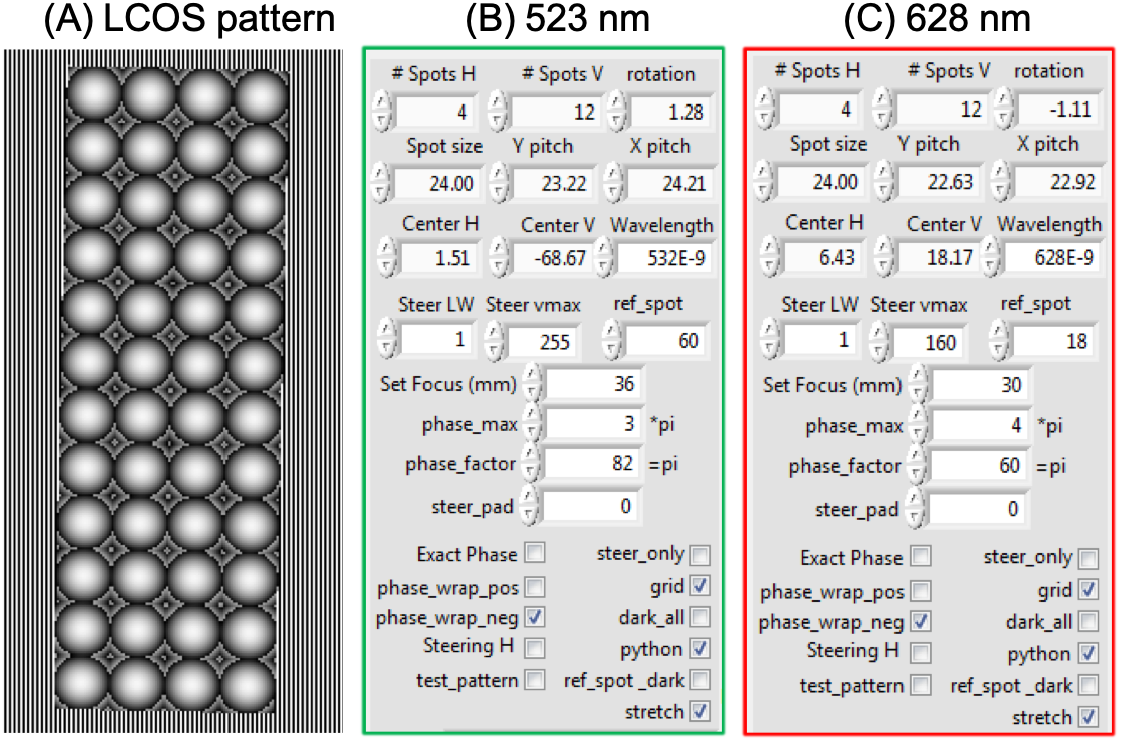
Pattern generation using two independent LCOS-SLMs. Focal lengths and beam-steering parameters differ for the two laser excitations. Adjustable parameters include number of spots, spot size, degree of rotation, pitch in X- and Y-directions, and pattern center (H, V) defined in LCOS-SLM units. During alignment, these parameters are optimized using the `LCOS_pattern_fitting notebook` (link). (A)LCOS-SLM generated 12×4 lenslet array surrounded by a periodic beam-steering pattern (shown for the green laser only). (B) and (C) show experimentally derived LCOS-SLM parameters for the spots and beam-steering patterns of the green (532 nm) and red lasers (628 nm) respectively.

Fluorescence emission is focused by the microscope tube lens, *L*_2_. The microscope’s internal flippable mirror, *M_I_* is used to toggle between the side and bottom ports of the microscope. A CMOS camera (Grasshopper3 GS3-U3-23S6M-C, FLIR, BC, Canada) is attached to the side port, and is used for alignment purposes. The bottom port directs the emission fluorescence to a recollimating lens, *L*_4_ (*f* = 100 mm, AC254-100-A, Thorlabs). Light is then split with an emission dichroic mirror, *DM_EM_* (Brightline Di02-R635, Semrock), spectral leakage from the red laser and Raman scattering due to the green laser being filtered on the donor emission path by an additional band-pass filters (donor: Brightline FF01-582/75, Semrock).

Each signal is focused on its respective SPAD array by lens *L*_5_ (*f* = 150 mm, AC254-150-A, Thorlabs). Each SPAD array is mounted on micro-positioning stage allowing adjustments of the detectors in all three dimensions. The detectors can be precisely aligned in the x and y directions using software controlled open-loop piezo-actuators (P/N 8302; drivers: P/N 8752 and 8753; Newport Corporation, Irvine, CA).

Each SPAD array is equipped with a field-programable gate array (FPGA; Xilinx Spartan 6, model SLX150), a humidity sensor, and a USB connection for monitoring time-binned counts and humidity levels. The FPGA provides 48 parallel and independent streams of LVDL pulses, which are converted to TTL pulses before they are fed to a programmable counting board (PXI-7813R, National Instruments, Austin, TX) providing 12.5 ns resolution time-stamping and a channel ID for each pulse. The LabVIEW code programming the FPGA module is available in the `Multichannel-Timestamper` online repository (link). Note that the specific acquisition board used in this work is not in production anymore, but can be found online from third party vendors. An alternative is one of the PXI-78XYR boards (X=3,4,5; Y=1,2,3,4) which provide 96 digital inputs and higher performance FPGAs.In this section, we provide an outline of the different steps involved in a typical multispot analysis workflow. Details can be found in previous publications and their associated Supporting Information files [63, 29, 30, 54].

## Appendix C Data Analysis

In this section, we provide an outline of the different steps involved in a typical multispot analysis workflow. Details can be found in previous publications and their associated Supporting Information files [63, 29, 30, 54].

### C.1 Photon streams

Photon streams are defined by the detection channel (D or A) and excitation period (for μsALEX: D or A, for PAX: D or *D* & *A*). Each photon is allocated to a stream based on its timestamp, *t_i_*, and its location, modulo the alternation period *T*, within the period, 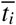 (Eq. 5, corrected for a possible offset, *t*_0_:

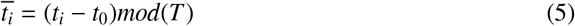

Because the transition between D-only to A-only or *D* & *A* excitation (and reciprocally) is not instantaneous due to the finite response time of the AOM (few *μs*), photons located within these transition periods are usually ignored due to their ambiguous origin [29]. They usually represent a small fraction of the total number of photons (< 5 %).

The histograms of 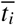 for the donor and acceptor channels are convenient to graphically define these ‘‘excitation periods” [29]. Table 1 indicates the notation used for the four photon streams in the two excitation periods. In μsALEX, both donor and acceptor channel histograms show large numbers of photons, while during the acceptor excitation period, only the acceptor channel histogram has a significant number of photon (the donor channel is limited to detector dark count). In PAX, the donor and acceptor channel histograms both contains significant numbers of photons during both D and DA (i.e *D* & *A*) excitation periods.

Due to this difference between μsALEXand PAX, a number of quantities defined in later sections take on different definitions.

**Table 1:** Photon streams for μsALEX and PAX alternation schemes. The excitation column indicates which laser is on during that period: D indicates 532 nm excitation and A indicates 628 nm excitation. Emission is detected in either the D or A channel.

| Alternation scheme | Excitation | Emission | photon streams |
| --- | --- | --- | --- |
| ALEX | D | D | $D_{ex}D_{em}$ |
| | D | A | $D_{ex}A_{em}$ |
| | A | A | $A_{ex}A_{em}$ |
| | A | D | $A_{ex}D_{em}$ |
| PAX | D | D | $D_{ex}D_{em}$ |
| | D | A | $D_{ex}A_{em}$ |
| | DA | D | $DA_{ex}D_{em}$ |
| | DA | A | $DA_{ex}A_{em}$ |

Raw photon streams denoted as, *F_X_ex_Y_em__*, corresponding to X excitation in the Y emission channel, are background corrected by subtraction of the background rate, *b_X_ex_Y_em__*, averaged over the whole period, times the burst duration, Δ*T*:

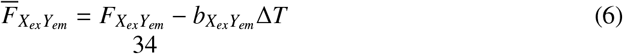

where

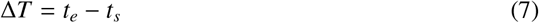

and *t_s_* (resp. *t_e_*) is the first (resp. last) timestamp in the burst.

In PAX, the (background corrected) total burst size is given by the sum of the background corrected photon streams (a similar definition holds in μsALEX, with *DA_ex_* replaced by *A_ex_*):

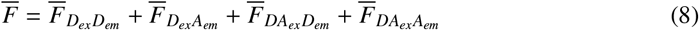

For FRET efficiency calculation, the total (corrected) fluorescence during donor excitation, 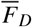, is used:

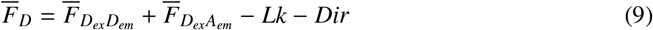

where *Lk* is the spectral leakage of the donor signal in the acceptor channel and *Dir* is the contribution of direct excitation of the acceptor dye by the green laser. The correction factors used to compute these quantities are discussed in section C.8.

In PAX, the 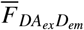 photon stream also contributes information, resulting in improved photon counting statistics compared to μsALEX. The PAX-specific definition of the corrected fluorescence emission during donor excitation is given by:

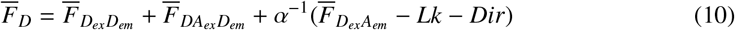

where *α* is defined as 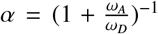, and *ω_A_* and *ω_D_* are the durations of the *DA_ex_* and *D_ex_* PAX alternation cycles, respectively. Typically the alternation periods have a duty cycle = 0.5 and *ω_A_*/*ω_D_* = 1. Multiplying by *α*^−1^ accounts for the continuous D-excitation by amplifying the 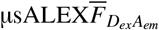 signal.

### C.2 Background rate estimation

Sources of background signal in single-molecule fluorescence experiments are due predominately to Raleigh and Raman scattering, scattered or out-of-focus fluorescence, the presence of sample or buffer impurities, and detector noise from DCR, crosstalk, or afterpulsing effects. Raleigh and Raman scattering can be effectively rejected by appropriate optical filters. Sample impurities cannot be totally eliminated, however, using spectroscopic grade reagents and buffer filtering greatly helps to reduce them.

Estimation of the background rate requires careful consideration. Rather than measuring a buffer only sample to use it as background, the background rate must be calculated for each measurement to account for scattering, out-of-focus fluorescent molecules and possible fluctuations during the measurement. One approach to estimating the background rate is to compute the inter-photon delay distribution, *φ*(*τ*), of each photon stream. The exponential inter-photon delay distribution for a Poisson process can be expressed as a weighted sum [88]:

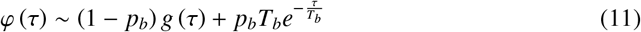

where *g*(*τ*) ∝ *τ*^−3/2^ is the distribution of inter-photon delays for a freely diffusing single-molecule in a Gaussian excitation volume and *T_b_* is equal to the average time between bursts [88]. The last term of Eq. 11 simply states that the background due to out-of-focus molecules can be described as a Poisson process with rate *b* = *T_b_*^−1^ (proportional to the concentration). The exponential term of the weighted sum dominates at long time-scales and is used to compute the background corrected inter-photon delay distribution.The background rate can for instance be estimated using the maximum likelihood estimator (MLE) for an exponential distribution:

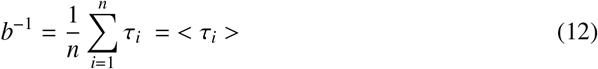

where the *τ_i_*’s are inter-photon delay times. Alternative estimators may be used, including the minimum variance unbiased estimator (MVUE) or the least-squared difference [29]. However, since only the long time-scale term in the inter-photon delay distribution is exponential, the background rate needs to be estimated using the exponential portion of *φ*(*τ*). The MLE of the restricted exponential distribution where *τ_i_* > *τ_min_* defines the background rate as:

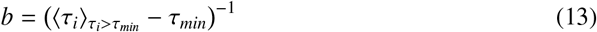

The choice for *τ_min_* is a compromise between estimation accuracy and data loss. A large *τ_min_* can result in a severely truncated data set giving unreliable statistics. Alternatively, a small *τ_min_* results in biased collection of short inter-photon delay times which are associated with singlemolecule diffusing within the center of the excitation PSF.

Determining an optimal *τ_min_* can be done automatically as discussed in [29].

Finally, in many smFRET experiments, the background rate may change over time, most commonly due to drift or evaporation, but possibly because of planned sample modifications. In the case of fluctuating backgrounds, the background rate estimation must be performed piecewise over time windows during which the rate is relatively constant (for rate estimation on the 48-spot setup we use a time window of 10 s).

In the case of multispot acquisition, these rate estimations need of course to be repeated for each spot.

### C.3 Burst search

After photon streams definition and background rates determination, the next step in smFRET analysis consists in a burst search, where fluorescent bursts due to single-molecules passing through the confocal volume are detected as “spikes” above the background signal. This is achieved with a ‘sliding window’ algorithm, first introduced by Seidel and collaborators [89, 57]. In each ‘sliding window’ of *m* sequential photons, the average photon (count) rate in one or more, or a sum of several photon streams, is calculated using the following definition:

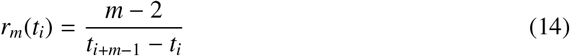

where *t_i_* the first time stamp of the series of *m* photons used to compute the rate [29]. A burst is identified if the count rate in that window is greater than a specified threshold rate. Typical values of *m* = 5 – 15 photons are used. Note that *m* also sets the minimum burst size.

Two methods can be used to specify the threshold rate:

1. a constant threshold can be set, or
2. an adaptive moving threshold can be used.

Using an adaptive threshold seamlessly takes account of possible background variations over time if the threshold is defined as proportional (factor *F*) to the local background rate. Typical values (*F* = 5 – 10) are generally appropriate and set the minimal signal-to-background rate (SBR) as (*F* – 1) [90]. A comparison of the choice of background threshold is presented in Ref. [29].

Typical burst searches are:

- “all photon burst search” (APBS): the burst search is performed the sum of all photon streams [60, 89, 57].
- “dual-channel burst search” (DCBS): two separate donor-only and acceptor-only burst searches are performed, and only bursts detected in both searches (and then only their common bursts) are retained [60].

The DCBS is useful for rejecting donor-only and acceptor-only species. In addition, by rejecting non-overlapping portions of D- and A-only bursts, the DCBS helps reducing the influence of photophysical effects such as blinking. Other burst searches may also be implemented. For example, the donor/acceptor emission burst search, *D_em_BS* or *A_em_BS*, selects all photons received in the donor or acceptor channel respectively, regardless of the laser alternation cycle. Similarly, the donor/acceptor excitation burst search, *D_ex_BS* or *A_ex_BS*, selects all photons received in the either channel during the D or A laser excitation period.

Both FRETBursts and ALiX allow burst searches to be implemented on arbitrary logical combinations of photon streams. While many options are available, it is often useful to begin an analysis using the APBS followed by burst selection (discussed in the next section). In this work, burst searches performed for multispot data were done independently for each spot, using a constant burst selection threshold on all photons (APBS), followed by further selections. A thorough evaluation of the effect of various burst searches on burst statistics is presented in Ref. [29].

### C.4 Fusing bursts

During analysis of freely diffusing molecules, it can be useful to “fuse” bursts separated by less than a specified minimum time, which typically correspond to the same molecule successively getting in and out of the excitation/detection volume. Fusing bursts results in bursts with more photons and, in general, better statistics, but assumes that no changes occur to the molecule in between crossing. This of course is not always the case [91]. However, fusing bursts with too long a minimum burst interval will increase noise due to additional background variance.

### C.5 Burst selection

A burst selection generally needs to follow the burst search, as it typically returns a large number of very small bursts contributing a large relative variance to any final burst statistics. Typically, a burst size selection is used that rejects bursts whose total size (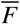, defined above in Eq. 8) falls below a set threshold (e.g. 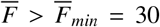 photons). In case different species are present in solution, selection needs to be performed after the initial burst search and all possible corrections are applied, in order to minimize bias in the selection process.

Other selections can be performed for specific purposes. For instance, in PAX, an additional burst selection based on the *DA_ex_A_em_* photon stream can be used in order to keep only FRET species. Computational details for the FRET burst searches and subsequent burst selections can be found in the `48-spot-smFRET-PAX-analysis` repository (link) [30].

### C.6 FRET efficiency (E) and stoichiometry ratio (S)

The ratiometric definition of FRET efficiency depends on the technique used (or more precisely, on the available photon streams) and can be quite difficult to properly calculate. However, in most cases, an approximate value neglecting corrections for quantum yield, detection efficiencies, absorption cross-section, etc., the so-called proximity ratio, is sufficient for distinguishing between sub-species and quantifying changes. For the sake of concision, we will limit ourselves to that latter definition. Exact definitions can be found in ref. [38] in the case of μsALEX, and in ref. [30] in the case of PAX.

Using background corrected burst sizes, 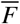, the proximity ratio, *E_PR_* can be expressed as:

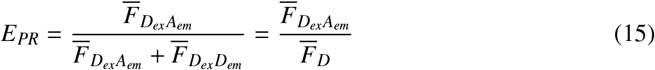

Where 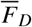 is the total background corrected fluorescence during donor excitation. The values of *E_PR_* range nominally from 0 to 1, where 0 indicates no FRET and 1 indicates 100% FRET, but because of imperfect background corrections, smaller and larger values are also possible.

Similarly, a fully corrected stoichiometry ratio, *S_γβ_*, can be defined in both μsALEX and PAX [38, 30], but the simpler uncorrected stoichiometry ratio, *S*, can be computed using only background-corrected burst sizes:

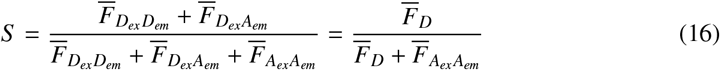

for μsALEX and:

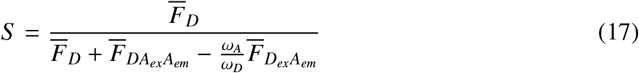

for PAX.

The stoichiometry ratio is used to separate donor-only species (i.e. singly-labeled molecules or doubly-labeled molecules with an inactive acceptor dye) and ranges nominally from 0 to 1, where *S* = 0 indicates acceptor-only species and *S* = 1 indicates donor-only species. Doubly-labeled molecules with active dyes, *i.e*. FRET species, are generally characterized by *S* ~ 1/2.

Note that the so-called unmodified stoichiometry ratio *S_u_* can be also used in PAX measurements:

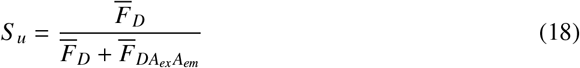

The benefit of *S_u_* over *S* is that *S_u_* results in a lower variance for small bursts, and thus can provide better separation between sub-populations. However *S_u_* depends on the FRET efficiency, namely *S_u_* decreases with increasing *E*, which could potentially impair sub-populations separation for low FRET efficiency species.

### C.7 E, S, and E – S Histograms

The 2-dimensional *E* – *S* histogram (or rather *E_PR_* – *S* (or *S_u_*) in the context of this discussion) allows separation of burst sub-populations according to their stoichiometry (S), and when relevant (doubly-labeled molecules) their proximity ratio (loosely speaking, according to their FRET efficiency or inter-dye distance). 1-dimensional projections along the *E_PR_* or *S* (or *S_u_*) direction, after selection of sub-populations of bursts, can be used to better visualize or quantify the distributions of *E_PR_* and *S* (or *S_u_*).

Quantitative analysis of these histograms is still a matter of debate, as burst search parameters affect these histograms in a complex manner. The most rigorous approach is one that uses information of each individual burst to compare observed and predicted histograms based on advanced modeling of the different experimental effects at play in the measurement (shot noise analysis [60, 29] or photon distribution analysis [92]). For a mere estimation of respective subpopulations and characteristic *E_PR_* or *S* (or *S_u_*) values for individual populations, fitting with an *ad-hoc* model qualitatively describing the observed histograms is appropriate.

Here, we use the following model of two asymmetric Gaussian distributions connected by a “bridge” corresponding to a sub-population of bursts due to coincident molecule detection, or bleaching/blinking events during transit:

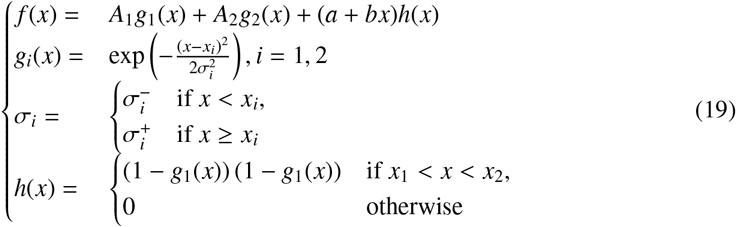

The integrals under each asymmetric Gaussian peak (*I_i_*) provide a good approximation of the number of bursts in each sub-population (without including the bridging bursts):

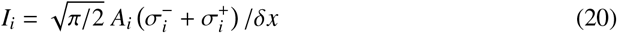

where *δx* is the histogram bin width.

### C.8 Correction factors

Accurate smFRET analysis requires the introduction of several correction factors *l, d, α, β*, and *γ*, using standard notations [38]. As mentioned previously, we will only discuss the first two for concision.

#### C.8.1 Donor leakage factor, l

The donor leakage factor, *l*, is defined via the relation:

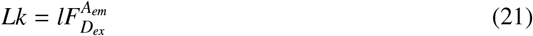

and can be expressed theoretically [38] in terms of *I_D_ex__*, the excitation intensity during the donor excitation period, 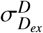, the absorption cross-section at the donor excitation laser wavelength, *ϕ_D_*, the quantum yield of the donor fluorophore, and 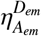, the donor emission detection efficiency in the acceptor channel.

The *l* correction factor is obtained experimentally from a donor-only (DO) histogram, by imposing that it is centered about 0 after correction. *l* can be calculated from a donor-only sample whose proximity ratio before correction is centered around *E_PR_DO__*, as:

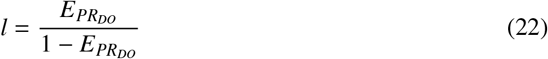

#### C.8.2 Direct acceptor excitation factor, d

The direct acceptor excitation correction factor, *d* is defined via the relation:

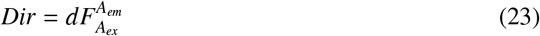

where *I_D_xe__* indicates the excitation intensity during the donor excitation cycle, 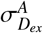 is the absorption cross-section of the acceptor dye under donor excitation, *ϕ_A_* is the quantum yield of the acceptor fluorophore, and 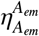 is the detection efficiency of acceptor emission in the acceptor channel.

*d* can be computed experimentally by imposing that the S histogram of an acceptor-only (AO) sample, be centered around 0 after correction. If *S_AO_* is the position of that histogram before correction:

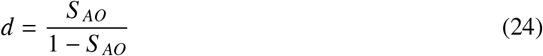

where *S_AO_* is the background corrected stoichiometry ratio (not corrected for *Lk* and *Dir*).

#### C.8.3 Other correction factors

As mentioned previously, other correction factors need to be introduced to compute accurate FRET efficiencies of stoichiometry ratios. Like *l* and *d* they in principle depend on the spot considered, and indeed, some, such as the *γ* factor, equal to the product of the A to D ratio of quantum yields and detection efficiencies, can be expected to be even more spot-dependent than *l* and d, due to differences of setup alignment in separate regions of field of view. However, provided that alignment is carefully done, we found out experimentally that spot-specific correction factor determination and inclusion does not significantly improve the separation of FRET subspecies [30].

### C.9 Burst statistics

Burst analysis can be used to quantify *E* and *S*, as well as other quantities related to concentration, diffusivity, brightness, etc. The following subsections will describe the statistics used in this article.

#### C.9.1 Burst Size

Burst size has been previously discussed in the context of burst selection. It is a useful quantity to histogram as it provides a quick preview of the data quality, small average burst sizes resulting in larger variance in any derived quantity. In the case of multispot data acquisition, the raw output of such an analysis is a series of similar (if not identical) size histograms, such as those shown in Fig. 14.

**Figure 14:**
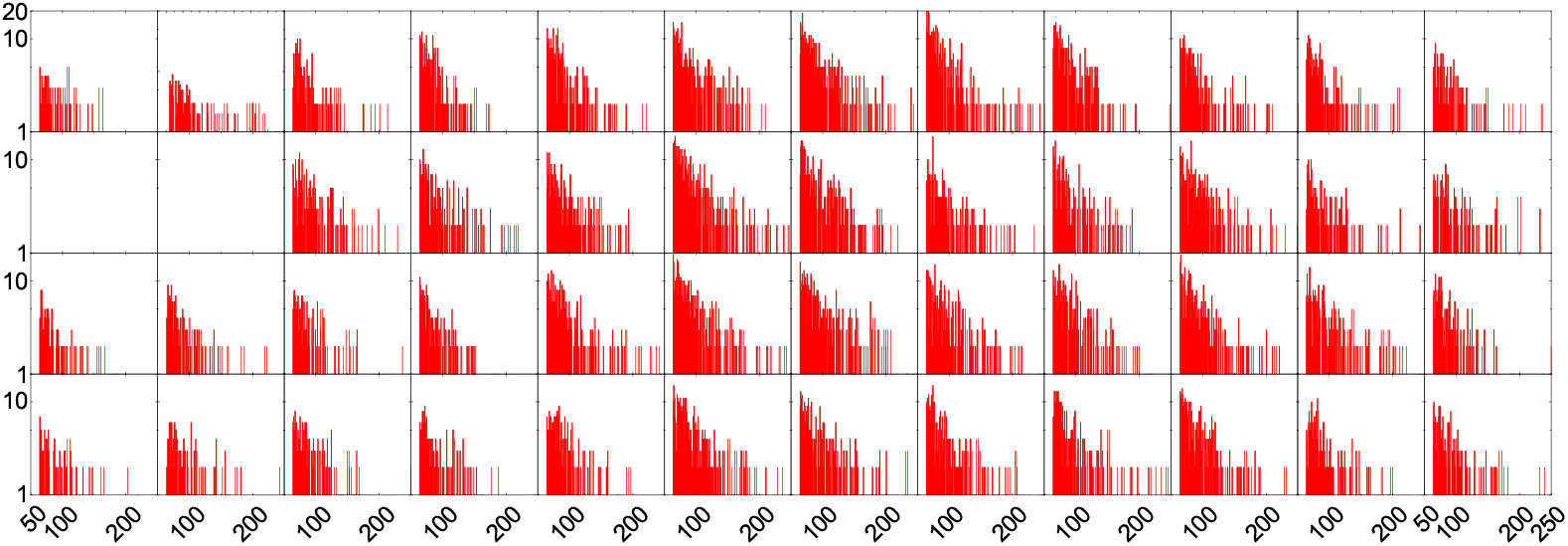
Burst size histograms (all photons stream) for each spot in the HT-smFRET microfluidic experiment discussed in Section 4.3.4. Analysis parameters: APBS, *m* = 10, *r_m_* ≥ 80 *kHz*, 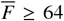. Spot 1 is at the top left, spot 12 at the top right. Spot 13 and 14 are missing from this series, due to a malfunction of two SPADs in the donor SPAD array. The better illumination of the center spots translates into larger burst statistics. Details of the analysis can be found in `ALiX Notebook, Flow, APBS, m= 10, Rmin = 80 kHz, Smin = 64.rtf` and associated files in the Figshare repository [81]).

When spot characteristics are similar, it is justified to pool these data into a single histogram, as shown in Fig. 15, for comparison between dataset acquired in the same conditions, or to assess the effect of different burst search parameters on the burst size distribution.

**Figure 15:**
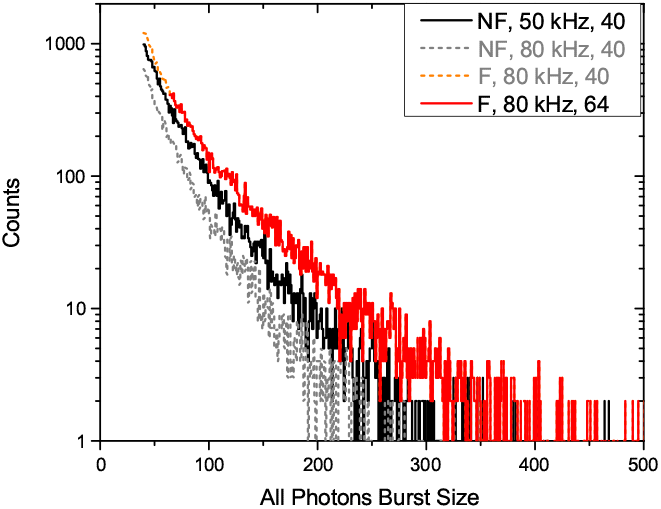
Pooled burst size (all photons) histograms corresponding to the two datasets discussed in section 4.3.4. The diffusion only (no flow or NF) dataset, recorded with lower excitation powers (by a factor ~ 1.6), was analyzed with a lower rate threshold (*r_m_* ≥ 50 *kHz*) for burst search and a lower burst size threshold (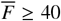, black) for burst selection, than the dataset recorded with flow (F, red), for which *r_m_* ≥ 80 *kHz* = 50 × 1.6, 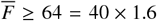, in order to obtain comparable number of bursts for analysis. For comparison, burst size distributions obtained when using the larger rate threshold for the no flow sample (*r_m_* ≥ 80 *kHz*, NF, gray), or the lower burst size threshold for the sample with flow (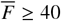, F, orange) are represented as dashed curves. The red curve corresponds to the sum of all histograms in Figure 14. The higher excitation powers used in the flow measurement more than compensate for the shorter transit time of molecules and more stringent burst search and selection criteria, as can be seen from the larger number and larger sizes of the collected bursts. Details of the analysis can be found in the different notebooks: `ALiX Notebook, XX, APBS, m = 10, Rmin = YY kHz, Smin = ZZ.rtf` where XX = Flow or No Flow, YY = 50 or 80, ZZ = 40 or 64, and associated files in the Figshare repository [81]).

#### C.9.2 Burst Duration

Burst duration has already been discussed in the context of burst search. Like burst sizes, it is an useful quantity to histogram for a quick overview of possible differences in spot sizes or alignment. Indeed, since the same sample is observed in all spots, the only expected scaling in case of similar spots, is a difference in the number of bursts (for instance if the excitation power is not uniform throughout the pattern). The overall shape of the duration histograms should in this case be identical, provided the proper burst search (constant threshold) is performed [29]. If burst duration histograms are dissimilar, sources of non-uniformities need to be investigated.

The burst duration distribution (and burst separation distribution) is however a complex function for which no analytical model currently exists. As discussed previously [29], a convenient representation of these complex distributions is a modified semilog histogram introduced by Sigworth & Sine [93] to study sums of exponentials, which has the advantage of allowing to easily identify the relevant time scale. In this “S&S” representation, data is binned logarithmically without normalization to account for the bins’ variable widths, and the square root of each bin content is displayed. An example of burst duration histograms obtained in the microfluidic HT-smFRET measurement discussed in section 4.3.4 is shown in Fig. 16.

**Figure 16:**
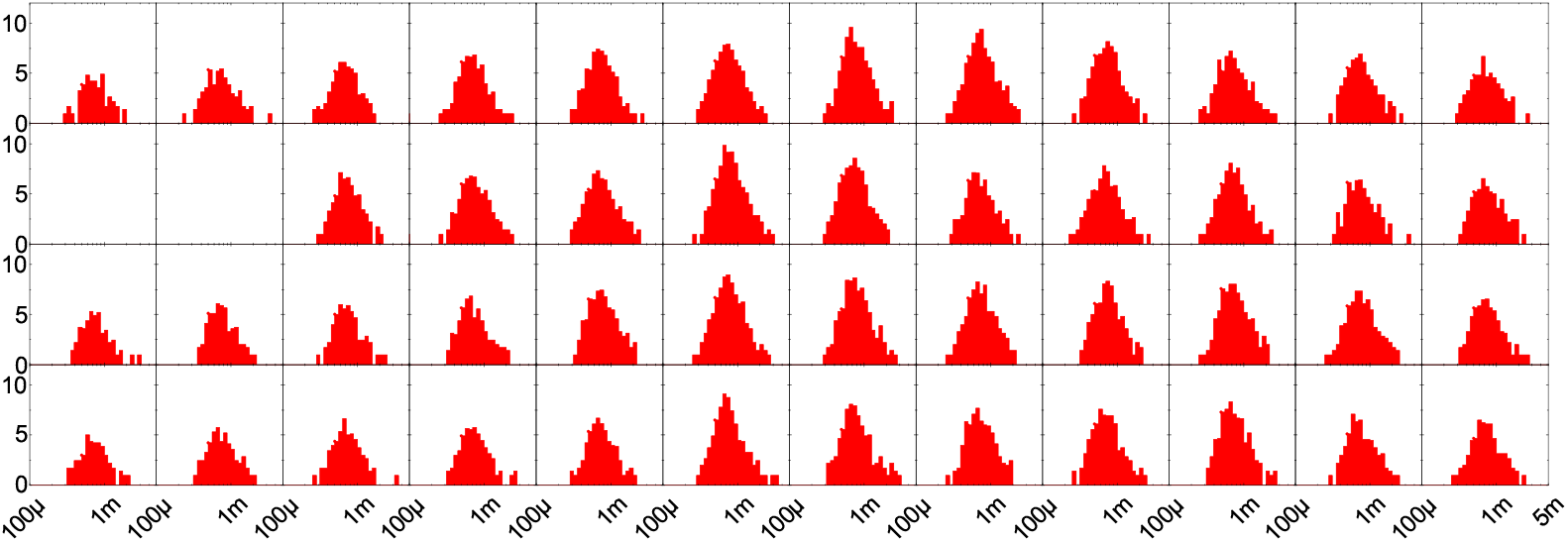
Burst duration histograms (unit: s) for each spot in the smFRET in flow experiment discussed in Section 4.3.4. Analysis parameters: APBS, *m* = 10, *r_m_* ≥ 80 *kHz*, 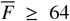. Spot 1 is at the top left, spot 12 at the top right. Spot 13 and 14 are missing from this series, due to a malfunction of two SPADs in the donor SPAD array. The better illumination of the center spots translates into larger burst statistics. Details of the analysis can be found in the notebook `ALiX Notebook, Flow, APBS, m = 10, Rmin = 80 kHz, Smin = 64.rtf` and associated files in the Figshare repository [81]).

As for burst sizes, if the spot parameters differ little, it is justified to pool these data into a single histogram, as done in Fig. 17 for comparison with data taken in the same conditions.

**Figure 17:**
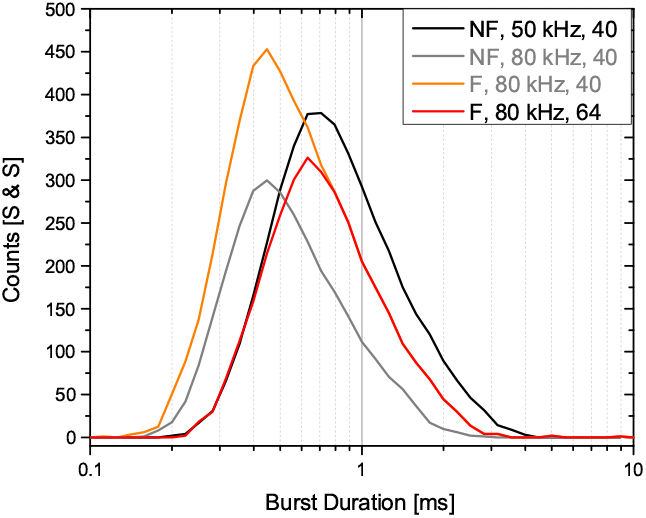
Pooled burst duration S & S histograms corresponding to the two datasets discussed in section 4.3.4. The diffusion only (no flow or NF) dataset, recorded with lower excitation powers (by a factor ~ 1.6), was analyzed with a lower rate threshold (*r_m_* ≥ 50 *kHz*) for burst search and a lower burst size threshold (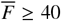, black) for burst selection, than the dataset recorded with flow (F, red), for which *r_m_* ≥ 80 *kHz* = 50 × 1.6, 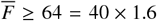, in order to obtain comparable number of bursts for analysis. For comparison, burst durations obtained when using the larger rate threshold for the no flow sample (*r_m_* ≥ 80 *kHz*, NF, gray), or the lower burst size threshold for the sample with flow (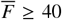, F, orange) are represented as well. The red curve corresponds to the sum of all histograms in Figure 16. The different burst search and selection criteria for each experiment result in different burst duration distributions, illustrating the challenges associated with this type of analysis. Details of the analysis can be found in the different notebooks: `ALiX Notebook, XX, APBS, m = 10, Rmin = YY kHz, Smin = ZZ.rtf` where XX = Flow or No Flow, YY = 50 or 80, ZZ = 40 or 64, and associated files in the Figshare repository [81]).

#### C.9.3 Peak Burst Count Rate

Due to diffusion of single molecules in the confocal excitation volume, burst quantities such as those discussed above are defined by probability densities which can sometimes be theoretically modeled [88], and in the most favorable cases, are asymptotically exponential. However, the choice of burst search parameters (photon stream, *m*, fixed or adaptive threshold, burst fusion, etc.), can affect the observed burst statistics. For example, applying a higher threshold to a burst that begins and ends with low count rates will result in a truncated burst that begins and ends earlier (the burst duration is decreased) and therefore has fewer photons (the burst size is reduced).

On the other hand, the peak count rate in a burst (maximum rate of photon detection defined using a particular number of photons) is usually obtained inside the burst (rather than at its edges) and therefore should not be affected by burst truncation.

Therefore, while quantities such as the burst size are related to precise trajectory of the molecule through the excitation PSF, the peak count rate reports only on how close to the spot excitation peak the trajectory brought the molecule. Histogramming this quantity for all bursts will thus report more directly on each spot’s peak excitation intensity, an important information in the comparison between spots in a multispot setup.

The definition of the peak count rate adopted in the Supporting Information of ref. [29] is:

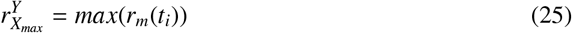

where the *t_i_*’s are timestamps within a burst and *r_m_*(*t_i_*) is defined by Eqn. 14.

The definition presented in Eqn. 25 does not account for laser alternation or which excitation cycle a timestamp arises from. To account for alternation, the peak count rate must be modified:

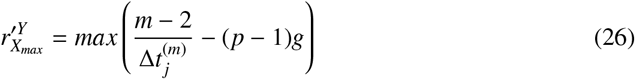

where the first and last timestamps of a burst are denoted as *t_j_* and *t*_*j*+*m*-1_. *g* is the minimum time between two donor excitation cycles, and *p* is the number of alternation periods separating the burst. As for the other statistics, the raw output of the analysis of a multispot dataset is a series of burst peak count rate histograms such as show in Fig. 18.

**Figure 18:**
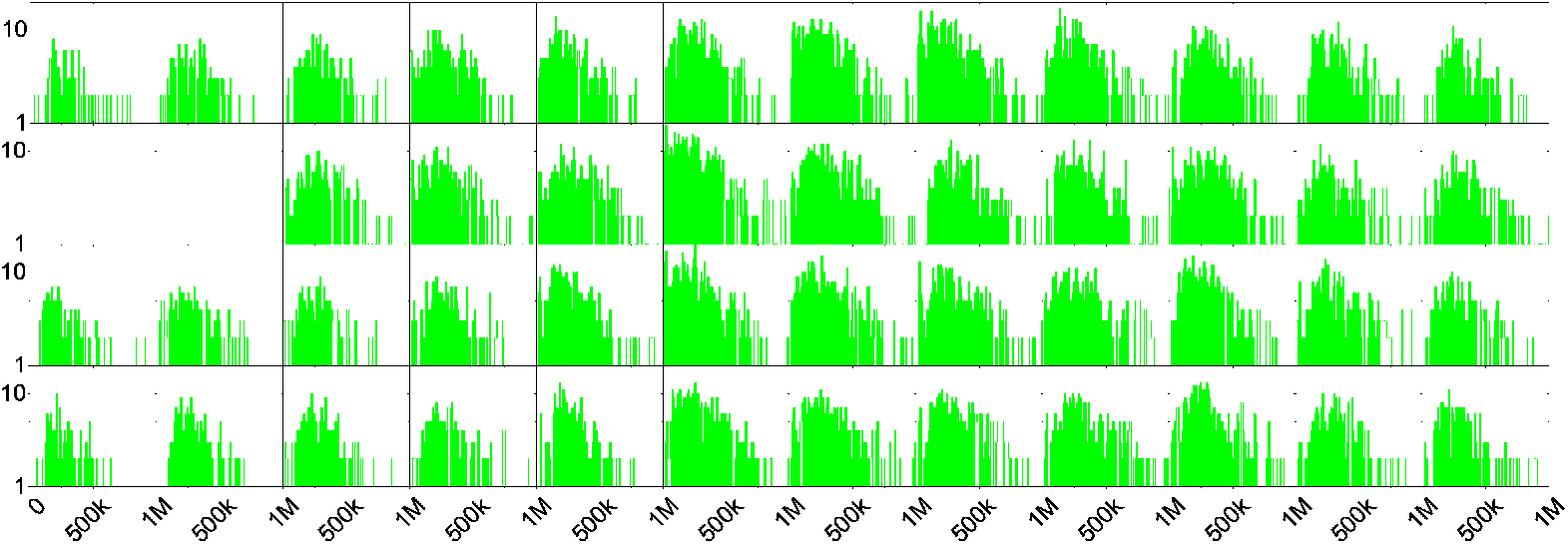
Burst peak count rate histograms during the D-excitation period and in the donor channel for each spot in the microfludic HT-smFRET experiment discussed in Section 4.3.4. Analysis parameters: APBS, *m* = 10, *r_m_* ≥ 80 *kHz*, 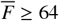. Spot 1 is at the top left, spot 12 at the top right. Spot 13 and 14 are missing from this series, due to a malfunction of two SPADs in the donor SPAD array. The better illumination of the center spots translates into larger number of bursts, but also larger peak burst rates. Details of the analysis can be found in the notebook `ALiX Notebook, Flow, APBS, m = 10, Rmin = 80 kHz, Smin = 64.rtf` and associated files in the Figshare repository [81]).

Some border spots clearly exhibit less and dimmer bursts, as could be expected from a close inspection of the spot intensity pattern shown in Fig. 12. Nevertheless, for a comparison of different experiments, pooling the burst peak count rates of all spots to form a single histogram is helpful, as done in Fig. 19.

**Figure 19:**
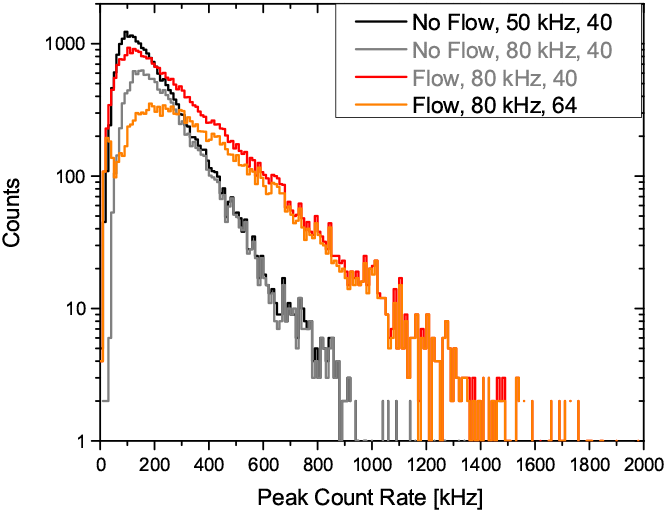
Pooled burst peak count rate histograms during the D-excitation period and in the donor channel corresponding to the two datasets discussed in section 4.3.4. The diffusion only (no flow or NF) dataset, recorded with lower excitation powers (by a factor ~ 1.6), was analyzed with a lower rate threshold (*r_m_* ≥ 50 *kHz*) for burst search and a lower burst size threshold (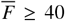, black) for burst selection, than the dataset recorded with flow (F, red), for which *r_m_* ≥ 80 *kHz* = 50 × 1.6, *F* ≥ 64 = 40 × 1.6, in order to obtain comparable number of bursts for analysis. For comparison, burst size distributions obtained when using the larger rate threshold for the no flow sample (*r_m_* ≥ 80 *kHz*, NF, gray), or the lower burst size threshold for the sample with flow (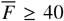, F, orange) are represented as dashed curves. The red curve corresponds to the sum of all histograms in Figure 18. As argued in the text, the asymptotic part of the burst peak count rate distribution is insensitive to the exact burst search and selection parameters used in the analysis, as is clear from the overlap of the exponential tails of the two no flow (NF, black and gray) and the two flow (F, red and orange) curves. The ratio of the two exponential coefficients (F: 216 kHz and NF: 116 kHz, *F/NF* = 1.9) is approximately equal to the ratio of the donor laser excitation powers used in the two measurements (500/300 = 1.7), as expected. Details of the analysis can be found in the different notebooks: `ALiX Notebook, XX, APBS, m = 10, Rmin = YY kHz, Smin = ZZ.rtf` where XX = Flow or No Flow, YY = 50 or 80, ZZ = 40 or 64, and associated files in the Figshare repository [81]).

### C.10 Fluorescence Correlation Analysis

Fluorescence correlation analysis (or spectroscopy, FCS) can be performed on single or multispot setups in order to characterize excitation/detection volumes sampled by the donor and acceptor, diffusion coefficients, brightness, and, provided enough statistics are available, short time-scale dynamics [46]. In the case of multispot experiments, FCS analysis is particularly helpful to detect otherwise difficult to quantify differences in spot characteristics, as the respective diffusion time through the excitation/detection volume, *τ_D_*, is one of the simplest pieces of information to extract from such an analysis and readily indicates differences between spots.

In past works, we have performed comparisons of single and multispot setups using FCS analysis [27, 29, 30]. Analysis may be performed on the same dye (autocorrelation function, ACF) or on two different dyes (cross-correlation function, CCF). This analysis has proven complementary to burst duration and brightness analysis, described in previous sections, to uncover subtle differences in effective excitation/detection volumes or peak excitation intensities [29].

However, quantitative FCS analysis suffers from many experimental artifacts and requires simplifying assumptions which are not always verified [94, 95]. In particular, current SPAD arrays suffer from measurable afterpulsing and additional effects at short time scales (< *μs*) which complicates the use of ACF as a routine tool.

CCF analysis on the other hand, eliminates most of these problems. In smFRET with two detection channels, it is limited to the correlation of donor and acceptor signals *within a spot*, but no such limitation exist when considering separate spots. In diffusion-only experiments, cross-correlating the signals of different spots does not provide much information (except for a measure of the optical crosstalk between pixels, if that analysis is performed within a single detection channel [29, 54]), because the distance between spots (~ 5 *μm*) is too large to extract any diffusion coefficient information.

However, as illustrated in Section 4.3.3, CCF analysis between SPADs from a single detection channel can be used to extract flow velocity (and direction, if needed be [79]). In particular, as for other multispot statistics, the average CCF of all spots can be analyzed for increased statistical accuracy, as done in Fig. 10.

Future multispot setups may involve two SPAD arrays per channel, allowing CCF analysis within single spots and channels, providing access to short timescale dynamics. When taking proper account of differences between spots, averaging of CCF curves from multiple spots could considerably decrease the time necessary to accumulate enough statistics for short time scale dynamics studies [96].

